# Using soybean historical field trial data to study genotype by environment variation and identify mega-environments with the integration of genetic and non-genetic factors

**DOI:** 10.1101/2022.04.11.487885

**Authors:** Matheus D Krause, Kaio O G Dias, Asheesh K Singh, William D Beavis

## Abstract

1

Soybean (*Glycine max* (L.) Merr.) provides plant-based protein for global food production and is extensively bred to create cultivars with greater productivity in distinct environments. Plant breeders evaluate new soybean genotypes using multi-environment trials (MET). The application of MET assumes that trial locations provide representative environmental conditions that cultivars are likely to encounter when grown by farmers. In addition, MET are important to depict the patterns of genotype by environment interactions (GEI). To evaluate GEI for soybean seed yield and identify mega-environments (ME), a retrospective analysis of 39,006 data points from experimental soybean genotypes evaluated in preliminary and uniform field trials conducted by public plant breeders from 1989-2019 was considered. ME were identified from phenotypic information from the annual trials, geographic, soil, and meteorological records at the trial locations. Results indicate that yield variation was mostly explained by location and location by year interactions. The static portion of the GEI represented 26.30% of the total yield variance. Estimates of variance components derived from linear mixed models demonstrated that the phenotypic variation due to genotype by location interaction effects was greater than genotype by year interaction effects. A trend analysis indicated a two-fold increase in the genotypic variance between 1989-1995 and 1996-2019. Furthermore, the heterogeneous estimates of genotypic, genotype by location, genotype by year, and genotype by location by year variances, were encapsulated by distinct probability distributions. The observed target population of environments can be divided into at least two and at most three ME, thereby suggesting improvements in the response to selection can be achieved when selecting directly for clustered (i.e., regions, ME) versus selecting across regions. Clusters obtained using phenotypic data, latitude, and soil variables plus elevation, were the most effective. In addition, we published the R package SoyURT which contains the data sets used in this work.

**Highlights:**

- Mega-environments can be identified with phenotypic, geographic, and meteorological data.
- Reliable estimates of variances can be obtained with proper analyses of historical data.
- Genotype by location was more important than genotype by year variation for seed yield.
- The trend in genotype by environment variances was captured in probability distributions.

## 4 Introduction

The terms genotype (G) and phenotype (P) were first coined by Johannsen (1911) after the rediscovery of Mendel’s work. Since then, the understanding of the mapping function that links G to P has been an on-going research interest (Pigliucci 2001, p. 2). The mapping of G to P for most quantitatively expressed traits is complicated by the differential response of genotype(s) to different environments, i.e. genotype by environment interactions (GEI), wherein phenotypic variation is shaped by G, Environment (E), and GEI (Tabery 2008; Sprague and Federer 1951). The GEI component will increase P variance and reduces heritability estimates, complicating breeding decisions and lowering response to selection. Additionally, it leads to variation in the adaptation and plasticity of genotypes’ response to targeted agroecological zones (Mackay *et al*. 2019; Cooper and DeLacy 1994; Haldane 1947). Hence, the GEI is particularly important to breeders as they explore the responsiveness of varieties for specific or broad environments.

Plant breeders evaluate candidate genotypes in multi-environment trials (MET) to reveal GEI patterns (Oakey *et al*. 2016; Smith *et al*. 2001b). MET utilize locations sampled from a target population of environments (TPE) representing farm production environments, i.e. the growing conditions that an experimental genotype is expected to encounter as a cultivar grown by farmers (Bustos-Korts *et al*. 2021). Hence, a TPE comprises many environments, spatially across agroecological zones and temporally over the years (Crespo-Herrera *et al*. 2021). The manifestation of GEI in a TPE has two components, the “static” environmental characteristics such as soil, longitude, latitude, and “non-static” seasonal environmental characteristics such as weather and management practices (Cullis *et al*. 2000). If GEI is large and associated with consistent sub-groupings of environments within the TPE, greater gains from selection should be achieved by subdividing locations into Mega-Environments (ME) (Crespo-Herrera *et al*. 2021; Yan 2016; Atlin *et al*. 2000a).

According to CIMMYT (1989, p. 58), “ME are broad, not necessarily contiguous areas, defined by similar biotic and abiotic stresses, cropping system requirements . . . ”. Another definition is a group of environments that share the same winning genotypes (Kang 2020; Gauch and Zobel 1997), or that within ME there is minimal crossover interaction among the genotypes across environments (Singh *et al*. 2021; Smith *et al*. 2021). It should be noted that the terms subregion, region/regional, subdivision, clusters, zones, agroclimatic, ecogeographic, and ME are sometimes interchangeably used in the literature. One way of exploring GEI is to divide the TPE into ME, and to select within ME (Yan 2016). Some studies have investigated strategies to subdivide the TPE in maize (Windhausen *et al*. 2012), barley (Atlin *et al*. 2000b), wheat (George and Lundy 2019; Bustos-Korts 2017), sorghum (da Silva *et al*. 2021), alfalfa (Annicchiarico 2021), rice (Krishnamurthy *et al*. 2017), oat (Yan *et al*. 2010), and soybean (Zdziarski *et al*. 2019; Yan and Rajcan 2002).

There are several methods for dividing the TPE into ME. For example, the genotype main effect plus GEI (GGE) biplots (Yan *et al*. 2000) on soybean MET data was used by Zdziarski *et al*. (2019) to identify two ME in Midwestern Brazil with contrasting altitudes, levels of fertilizer, and incidence of soybean cyst nematode profiles. da Silva *et al*. (2021) and Krishnamurthy *et al*. (2017) also took advantage of GGE biplots to pinpoint ME for pre-commercial sorghum hybrids in Brazil and rice genotypes in India, respectively. For wheat, Crespo-Herrera *et al*. (2021) defined three ME in India with climate and soil data through principal component analysis, followed by a hierarchical clustering based on Euclidean distance with Ward’s method. For maize in Africa (CIMMYT’s program), Windhausen *et al*. (2012) explored historical (2001-2009) MET data to determine ME according to five subdivision systems (climate, altitude, geographic, country, and yield level), and concluded there was enough genotype by subregion interaction relative to genotypic variance to justify the selection for the low and high-yielding sub-regions separately. Other methodologies such as the additive main effects and multiplicative interaction (AMMI) model (Bustos-Korts 2017; Gauch and Zobel 1997) and factor analytic (FA) models (Smith *et al*. 2021; Bustos-Korts 2017; Smith *et al*. 2015, 2001b; Piepho 1997) also have been used. When ME are included in the model, it is called a zone-based model; therefore, yielding zone-based predictions (Buntaran *et al*. 2019). One of the main advantages of modeling ME in a mixed model framework is the ability to borrow information between zones from the genotype by ME interaction. This is particularly beneficial when fewer testing locations are available creating a sparse representation of genotypes in some ME (Piepho *et al*. 2016; Piepho and MOhring 2005).

The effectiveness of subdividing the TPE into ME was assessed by Atlin *et al*. (2000a) based on the theory of correlated response to selection, first applied to the GEI problem by Falconer (1952). Effective selection occurs when subdivision increases response to selection, which might occur if the genotype by ME interaction variance, i.e. genotype by region 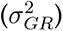, is large relative to the genotypic variance 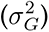. In terms of variance components derived from linear models, the GEI is composed of genotype by location 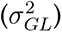, genotype by year 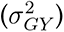, and genotype by location by year 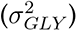 interaction variances. Both 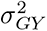 and 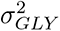 are non-static (unrepeatable) sources of variation. ME can be identified with the static portion of the 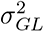, which is partially repeat-able across years (Yan 2016). When ME are identified and modeled, the 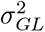 can be partitioned into 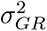 and genotype by location within ME 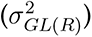. Furthermore, the 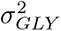 can be partitioned into a genotype by ME by year interaction 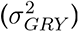, and genotype by location within ME by year 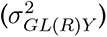 interaction (Atlin *et al*. 2000a). Consequently, accurate estimates of variance components are critical to properly identify ME.

Variance components can be estimated with unbalanced historical data to provide information for designing novel breeding strategies and optimizing resource allocation (Aguate *et al*. 2019). Efforts have been made to quantify component variability using historical MET data in wheat, maize, sunflower, sugar beet, potato, rye (Meyer *et al*. 2011; Laidig *et al*. 2008), among other commercial crops. However, proper modeling of historical data can be a significant challenge (Dias *et al*. 2020), and if not done properly can lead to misleading interpretations. In terms of variance component estimators, recent work from Aguate *et al*. (2019) and Hartung and Piepho (2021) considered both the imbalance of data (due to selection) and the properties of the residual maximum likelihood (REML) method (Patterson and Thompson 1971) to shed light onto potential bias of estimated variance components from MET using linear mixed models. Their results served as guidelines for designing the variance estimation portion of this work, which will be discussed later.

To identify and describe ME for soybean seed yield in the primary production area of North America, we obtained historical soybean performance data from Uniform Soybean Cooperative Tests (USDA 2021). We purposely chose this dataset because these trials have been used for decisions on cultivar release by public breeding organizations. The dataset consisted of 39,006 data points for maturity groups (MG) II and III spanning 63 locations between 1989 and 2019. Note that experimental genotypes were not evaluated at all locations within years and most were not evaluated for more than one year. The objectives of this study were to (*i*) investigate if the observed TPE spanning 31 years of field trial evaluations can be classified into ME, and (*ii*) estimate probability density functions for the underlying trend of genotypic, genotype by location, genotype by year, genotype by location by year, and residual variance components derived from linear mixed models. This modeling approach allowed us to fit parametric probability distributions to variance components to capture the GEI trend that can be used in future simulation studies, needed for predicting plant breeding outcomes in changing climates. Currently, simulation studies rely on point estimates of variance components (Kleinknecht *et al*. 2016), or set heritability values such as low or high (Rutkoski 2019). By capturing the GEI trend using historical data, we generate reliable variance estimates that can be used to conduct more realistic simulation studies.

## 5 Data and Methods

### 5.1 Phenotypic data

Annual PDF reports from the Northern Region of the Uniform Soybean Tests were obtained from the USDA-ARS (see Section 12). Information from the PDF files was transcribed into CSV format files. The retrieved data for MG II and III contain the empirical best linear unbiased estimated values (eBLUE) of genotypic means for seed yield adjusted to 13% moisture and reported in bushels per acre (bu/ac). Information from the coefficient of variation (CV%), genotypic variance (see Section 5.3.1), and the number of blocks per trial was also obtained. Before data analyses, individual trials with estimates of reliability (*i*^2^, see Section 5.3.1) lower than 0.10, coefficients of variation (CV%) greater than 20%, and individual observations lower than ten bu/ac were removed. A full description of the data cleaning is given in Appendix A.

The trials were divided into MG, Preliminary (PYT), and Uniform Regional (URT) Yield Trials. Because there are large numbers of experimental genotypes created by several public breeding programs within each MZ, the PYT are further split into two groups: PYT-A and PYT-B. Additionally, experimental genotypes with introgressed transgenic alleles were evaluated independently in trials referred to as PYT/URT-RR or PYT/URT-TM, depending on whether the lines included Roundup Ready or other transgenic traits. Preliminary analyses revealed that estimates of variance components in MG II were not significantly different than in MG III. Thus, subsequent analyses were conducted on combined data from both MG (Appendix B).

Operationally, experimental genotypes were first evaluated in PYT, and if not culled, were subsequently evaluated in URT. Most experimental genotypes are culled on an annual basis, thus only 0.47% of all potential combinations of experimental genotypes, locations, and years were evaluated. Field trials were laid out in randomized complete blocks (RCBD) with two or more blocks. In a given year, a PYT was usually conducted at eight or more locations. Experimental genotypes retained for regional trials were evaluated at 12 or more locations in URT. Some experimental genotypes might be evaluated in two subsequent years of URT. In addition to the experimental genotypes, entries in each field block included common check varieties (*∼* 3). We noted that check varieties were seldom retained for more than four consecutive years. For more information about the trial field plot design and agronomic practices, please refer to the PDF files.

### 5.2 Environmental data

In addition to phenotypic (PHE) data from yield trials, environmental data associated with trial locations were obtained. Elevation information was obtained from the “elevatr” package (Hol- lister *et al*. 2021). Soil characteristics at a depth of 5-15 cm were downloaded from Soilgrids (https://soilgrids.org/) with a modified R script available at https://github.com/zecojls/downloadSoilGridsV2, and further processed with the package “raster” (Hijmans 2021). The soil characteristics are referred to as soil variables (SV) and include bulk density (SV1), cation exchange capacity (SV2), clay content (SV3), total nitrogen content (SV4), pH (SV5), sand content (SV6), silt content (SV7), and organic carbon content (SV8). Detailed information about SV are available on the Soilgrids website. Latitudes for locations in the USA were downloaded from https://simplemaps.com/data/us-cities, and Canadian locations were obtained using Google Maps. Meteorological data, referred herein as MV for each location were obtained from “NASA’s Prediction of Worldwide Energy Resources” (NASA POWER, https://power.larc.nasa.gov/) with the package “nasapower” (Sparks 2018), and further processed with the “EnvRtype” package (Costa-Neto *et al*. 2021). In total, 19 MV were retrieved daily (averages) from the average planting date until the average check variety maturity date (R8) for each environment (location by year combination). A summary of the environmental variables is provided in Appendix C (Tables C1 and C2), and for more detailed information, please refer to the cited references.

**Table 1:** Models and criteria used to evaluate the models for purposes of clustering locations into groups representing the most likely target population of environments. The best-fit model (M2-18) is highlighted in bold.

| Model | Covariance structure<sup>a</sup> |  |  | Evaluation criteria |  |  |
| --- | --- | --- | --- | --- | --- | --- |
| | $\Sigma_{gl}$ | $\Sigma_{gy}$ | $\Sigma_{\epsilon}$ | AIC | % Var(GL) | % Var(GY) |
| M2-1 | $\mathbf{I} \otimes \mathbf{I}$ | $\mathbf{I} \otimes \mathbf{I}$ | $\Sigma_3$ | 175294 | - | - |
| M2-2 | $\mathbf{D} \otimes \mathbf{I}$ | $\mathbf{I} \otimes \mathbf{I}$ | $\Sigma_3$ | 174256 | - | - |
| M2-3 | $\mathbf{I} \otimes \mathbf{I}$ | $\mathbf{D} \otimes \mathbf{I}$ | $\Sigma_3$ | 174964 | - | - |
| M2-4 | $\mathbf{D} \otimes \mathbf{I}$ | $\mathbf{D} \otimes \mathbf{I}$ | $\Sigma_3$ | 173951 | - | - |
| M2-5 | $\mathbf{FA}_1 \otimes \mathbf{I}$ | $\mathbf{D} \otimes \mathbf{I}$ | $\Sigma_3$ | 172845 | 48.5 | - |
| M2-6 | $\mathbf{FA}_2 \otimes \mathbf{I}$ | $\mathbf{D} \otimes \mathbf{I}$ | $\Sigma_3$ | 172560 | 67.3 | - |
| M2-7 | $\mathbf{FA}_3 \otimes \mathbf{I}$ | $\mathbf{D} \otimes \mathbf{I}$ | $\Sigma_3$ | 172359 | 78.2 | - |
| M2-8 | $\mathbf{FA}_4 \otimes \mathbf{I}$ | $\mathbf{D} \otimes \mathbf{I}$ | $\Sigma_3$ | 172303 | 87.6 | - |
| M2-9 | $\mathbf{FA}_1 \otimes \mathbf{I}$ | $\mathbf{FA}_1 \otimes \mathbf{I}$ | $\Sigma_3$ | 172754 | 48.0 | 63.4 |
| M2-10 | $\mathbf{FA}_2 \otimes \mathbf{I}$ | $\mathbf{FA}_2 \otimes \mathbf{I}$ | $\Sigma_3$ | 172472 | 66.2 | 82.5 |
| M2-11 | $\mathbf{FA}_3 \otimes \mathbf{I}$ | $\mathbf{FA}_3 \otimes \mathbf{I}$ | $\Sigma_3$ | 172288 | 77.0 | 96.4 |
| M2-12<sup>b</sup> | $\mathbf{FA}_4 \otimes \mathbf{I}$ | $\mathbf{FA}_4 \otimes \mathbf{I}$ | $\Sigma_3$ | - | - | - |
| M2-13 | $\mathbf{FA}_2 \otimes \mathbf{I}$ | $\mathbf{FA}_1 \otimes \mathbf{I}$ | $\Sigma_3$ | 172494 | 66.4 | 61.0 |
| M2-14 | $\mathbf{FA}_3 \otimes \mathbf{I}$ | $\mathbf{FA}_1 \otimes \mathbf{I}$ | $\Sigma_3$ | 172296 | 77.6 | 61.0 |
| M2-15 | $\mathbf{FA}_3 \otimes \mathbf{I}$ | $\mathbf{FA}_2 \otimes \mathbf{I}$ | $\Sigma_3$ | 172276 | 77.5 | 82.9 |
| M2-16 | $\mathbf{FA}_4 \otimes \mathbf{I}$ | $\mathbf{FA}_2 \otimes \mathbf{I}$ | $\Sigma_3$ | 172205 | 84.7 | 86.3 |
| M2-17 | $\mathbf{FA}_4 \otimes \mathbf{I}$ | $\mathbf{FA}_3 \otimes \mathbf{I}$ | $\Sigma_3$ | 172234 | 87.3 | 96.7 |
| **M2-18** | **$\mathbf{FA}_5 \otimes \mathbf{I}$** | **$\mathbf{FA}_2 \otimes \mathbf{I}$** | **$\Sigma_3$** | **171990** | **90.2** | **87.2** |
| M2-18 | $\mathbf{FA}_5 \otimes \mathbf{I}$ | $\mathbf{FA}_3 \otimes \mathbf{I}$ | $\Sigma_3$ | 172196 | 88.9 | 96.9 |
| M2-19<sup>b</sup> | $\mathbf{FA}_6 \otimes \mathbf{I}$ | $\mathbf{FA}_2 \otimes \mathbf{I}$ | $\Sigma_3$ | - | - | - |

<sup>a</sup> The evaluated variance-covariance structures were identity ( $\mathbf{I}$ ), diagonal ( $\mathbf{D}$ ), and factor analytic ( $\mathbf{FA}_k$ ) from order $k = 1, \dots, 6$ . $\Sigma_{gl}$ and $\Sigma_{gy}$ are covariance matrices for genotype by location (GL) and genotype by year (GY) interactions, respectively. $\Sigma_{\epsilon} = \Sigma_3$ is the residual variance matrix assumed to be known from single trial and location analysis.
<sup>b</sup> Singularity in the Average Information matrix.

### 5.3 Data analyses

A stage-wise approach was followed (Piepho *et al*. 2012; Smith *et al*. 2001a; Frensham *et al*. 1997). The first-stage model was applied to individual trials within locations resulting in the vector **y_1_** of eBLUE values, the second-stage across trials within locations (**y_2_**), and the third-stage across locations and years. All analyses were implemented using Asreml-R version 4 (Butler *et al*. 2017) in the R programming environment (R Core Team 2021). Variance components of linear mixed models were estimated with REML followed by estimation/prediction of the fixed and random effects in Henderson’s mixed models (Henderson 1949, 1950; Henderson *et al*. 1959). When possible, computation time was sped-up with parallel processing by applying the doParallel and foreach packages (Microsoft Corporation and Weston 2020a,b).

#### 5.3.1 First-stage analyses

The first-stage analyses were previously performed by the collaborators (public soybean breeders) before the data were submitted to the USDA-ARS for aggregating and reporting as PDF files. Individual trials within locations were analyzed using a linear model in which genotypes and blocks were considered fixed effects, yielding eBLUE values of genotypic means (**y_1_**, entry-mean basis). The standard error (SE) of genotypic means was estimated as 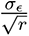, where *σ_E_* is the residual standard deviation calculated from the reported CV%, and *r* is the reported number of blocks for each trial. Because phenotypic values from each replicate are unavailable, we assumed all genotypes from a given trial had the same SE (i.e. equal replication, so that **y_1_** was obtained by ordinary least squares and not by generalized least squares). The *i*^2^ was then estimated as 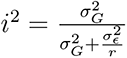, where 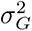 is the genotypic variance and 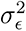 is the residual variance (Bernardo 2020, p. 173). The 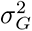 was estimated from the variance of entry means 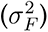, where 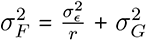.

#### 5.3.2 Second-stage analyses

The second-stage model embraced data from multiple trials with common entries within locations in a given year, as follows:

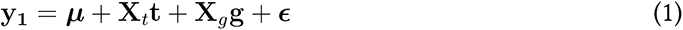

where **y_1_** is a vector of eBLUE values from the first-stage analyses, ***µ*** is the intercept, **X***_t_* is the incidence matrix of fixed effects of trials, ***t*** is a vector of fixed effects of trials, **X***_g_* is the incidence matrix of fixed effects of genotypes, ***g*** is a vector of fixed effects of genotypes, and ***E*** is a vector of residuals with ***E*** *∼* N(**0**, **Σ_1_**). The residual variance matrix **Σ_1_** is a diagonal matrix with elements equal to 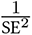 (Smith *et al*. 2001a; Frensham *et al*. 1997), where SE was estimated from the first-stage analyses. Estimated values from Model 1 (**y_2_** and SE) are available in the R package SoyURT.

#### 5.3.3 Multi-location and multi-year analyses

The third-stage analysis used the “baseline” three-way model for multiple locations and years:

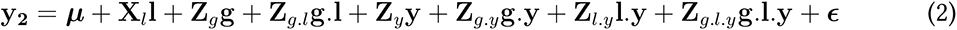

where **y_2_** is a vector of eBLUE values from the second-stage analyses, ***µ*** is the intercept, ***l*** is a vector of fixed effects of locations, ***g*** is a vector of random effects of genotypes with 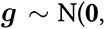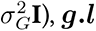 is a vector of random effects of genotype by location interactions with 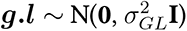, ***y*** is a vector of random effects of years with 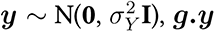 is a vector of random effects of genotype by year interaction with 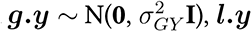 is a vector of random effects of location by year interaction with 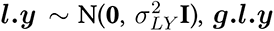 is a vector of random effects of genotype by location by year interaction with 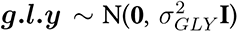, and ***E*** is a vector of residuals with ***E*** *∼*N(**0**, **Σ_2_**). **X***_l_*, **Z***_g_*, **Z***_g.l_*, **Z***_y_*, **Z***_g.y_*, **Z***_l.y_*, and **Z***_g.l.y_* are incidence matrices for their respective effects. The elements of the residual variance matrix **Σ_2_** were obtained from Model 1.

#### 5.3.4 Probability distributions of estimated variance components

A modified jackknife resampling approach was used to obtain empirical probability distributions for the variance components 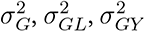, and 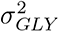. Utilizing results from Aguate *et al*. (2019) and Hartung and Piepho (2021), the data were divided into four groups representing consecutive eras of soybean cultivar development: From 1989 to 1995, from 1996 to 2003, from 2004 to 2011, and from 2012 to 2019. For the first group (1989-1995), there were 181 environments (location-year combination); for 1996-2003, 194 environments; for 2004-2011, 100 environments; and for 2012-2019, 116 environments. The modified jackknife approach consisted of leaving-one-environment out (instead of one observation), and then obtaining REML estimates of variance components with a modified version of Model 2 that considered locations as a random effect with variance 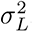. Estimates of variance components were combined and evaluated for a best- fit probability distribution with the package “ForestFit” (Teimouri 2021). Given the lack of data from individual plots, trial-based estimates of 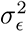 from the first-stage of analyses were used (i.e., no resampling). Distributional parameters were estimated via the expectation maximization (EM) algorithm (Dempster *et al*. 1977) using the log-likelihood functions of the Gamma, Log-Logistic, Log-Normal, Burr, and F univariate and multivariate distributions. In addition to a visual comparison of the modeled distributions relative to the empirical distributions, Akaike (AIC) and Bayesian (BIC) information criteria (Akaike 1974; Schwarz 1978) as well as the Kolmogorov-Smirnov (KS), Cramer-von Mises (CM), and Anderson-Darling (AD) goodness-of-fit statistics (Stephens 1986) were considered to select the best-fit distribution for each variance component. A classical penalized criteria based on the loglikelihood (AIC, BIC) provided protection from overfitting.

#### 5.3.5 Identification of mega-environments

Herein, “ME” and “cluster” are used interchangeably. We clustered 63 locations using six criteria: (i) phenotypes, i.e. seed yield (PHE), (ii) eight soil variables (SV) plus elevation (SoilE); (iii) latitude, where locations were split into two groups (Lat2), (iv) latitude, where locations were split into three groups (Lat3); (v) 19 meteorological variables (MV) with means across years (WA); and (vi) MV with means nested within years (WW).

Except for Lat2 and Lat3, the optimal number of clusters was then defined based on the Silhouette and Elbow methods using the package factoextra (Kassambara and Mundt 2020), followed by a K-means clustering with the R base function *kmeans()* allowing for a maximum of 1,000 iterations and 100 multiple initial configurations of the *K* groups.

#### 5.3.5.1 Clustering with PHE data

Several variations of Model 2 were evaluated by modeling the variance-covariance structures (VCOV) for the genotype by location (Σ*_gl_*) and genotype by year (Σ*_gy_*) interaction terms. The simplest model (M2-1, baseline) assumed independent effects for Σ*_gl_* and Σ*_gy_*. The next set of models allowed heterogeneous variances for Σ*_gl_* (M2-2), Σ*_gy_* (M2-3), or both (M2-4). Specific pairwise covariances for both Σ*_gl_* and Σ*_gy_* were assessed with models M2-5, M2-6, …, M2-20 (Table 1). The VCOV structures included identity (**I**), diagonal (**D**), and factor-analytic (**FA**(*k*)) of order *k* (Piepho 1997; Smith *et al*. 2001b, 2015). Models M2-5, M2-6, …, M2-20 are variations of the FA for different *k^t^s*.

In addition to the AIC selection criteria to select the best-fit model, the overall percentage of genetic variance accounted by each *k* factor in FA models was also considered. For each model term (Σ*_gl_*, Σ*_gy_*), it is defined as 100[tr(**ΛΛ***^t^*)/tr(**ΛΛ***^t^* + **Ψ**)], where “tr” is the trace of the matrix, **Λ** (63 *× k*) is the matrix of loadings, and **Ψ** (63 *×* 63) is a diagonal matrix of specific variances associated with each location. With the best-fit FA model, locations were clustered based on the estimated Σ*_gl_* loadings (Bustos-Korts 2017; BurgueNo *et al*. 2008) after Varimax rotation. Genetic correlations between locations (**C**) were further estimated by **C** = **DGD**, where **G** = (**ΛΛ***^t^* + **Ψ**) is the estimator of genetic variances, and **D** is a diagonal matrix composed by the inverse of the square root of the diagonal values of **G** (Smith *et al*. 2015).

#### 5.3.5.2 Clustering with SV data

First the SV (including elevation) were centered and scaled to unit variance. Subsequently, a principal component analysis (PCA) by non-linear iterative partial least squares (Wold 1966) was performed to reduce collinearity with the pcaMethods package (Stacklies *et al*. 2007). The number of principal components (PC) was selected with a 90% threshold of cumulative variance explained, followed by a Varimax rotation.

#### 5.3.5.3 Clustering with MV data

Meteorological data was retrieved for each environment for the growing season. Instead of using averaged values for each MV across the season, we used the Critical Environmental Regressor through Informed Search (CERIS) procedure proposed by Li *et al*. (2018) to identify relevant periods during growth and development. The method consists of screening meteorological data across environments to identify a period (window) of days after planting with the highest Pearson correlation between the population average (i.e. environmental average) and the MV. The idea is to identify periods of meteorological data most likely to affect growth and development associated with the phenotypic data (yield, plant height, etc.). Instead of using environmental averages (the average value of **y_2_** for each environment) to identify the window, we further modified CERIS to account for genotype by environment deviations within years, as follows:

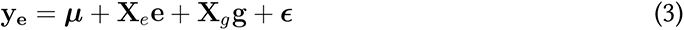

where **y_e_** is a vector of eBLUE values in a given year (i.e. **y_2_** in a given year), ***µ*** is the intercept, ***e*** is a vector of fixed effects of environments, ***g*** is a vector of fixed effects of genotypes, and ***E*** is a vector of residuals with ***E*** *∼* N(**0**, **Σ_e_**). **X***_e_* and **X***_g_* are incidence matrices for their respective effects, and **Σ_e_** is the residual variance matrix obtained from Model 1.

Each environment within years (i.e. a location) was then represented as the average of the squared residuals, which was used by CERIS. Model 3 was exclusively designed to replace environmental means with a metric more useful to identify ME with repeatable GEI. The residuals represent the GL deviations nested within years, which are confounded with GLY effects. An alternative model that contains the genotype by environment interaction is presented in Appendix D.

The CERIS was computed in two ways: across (WA) and within (WW) years, and the best window (i.e. highest correlation) for each MV was selected. In both cases, the final dataset used for clustering consisted of a matrix of 63 rows (locations) by 19 columns (MV). For WA, the algorithm (*i*) selected the best window across the 591 observed environments, (*ii*) the average value of each MV was computed with the selected window for each year a given location of observed, and (*iii*) locations were represented with the averaged value across years. For WW, the only modification is at step (*i*): each year a given location was observed, the algorithm selected its best window. A minimum window size of seven days was considered in all cases. After identifying the most relevant dataset of MV, the data were centered and scaled to unit variance. Subsequently, clustering was conducted as described for the SV.

#### 5.3.5.4 Effectiveness of clustering

We used the ratio of correlated responses from selection across all environments relative to direct responses to selection within ME (CR/DR) (Atlin *et al*. 2000a; Bustos-Korts 2017) as a metric to assess the relative effectiveness of clustering locations into ME. The zone-based model is defined as follows:

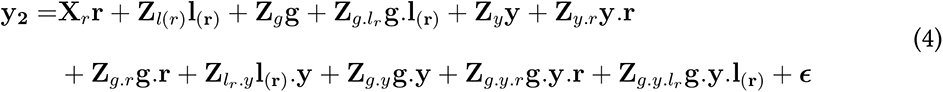

where ***r*** is a vector of fixed effects of clusters, and ***l*_(_*_r_*_)_**, ***g.l*_(_*_r_*_)_**, ***y.r***, ***g.r***, ***l_r_.y***, ***g.y.r***, and ***g.y.l_r_***, are random vectors with specific variances of locations within clusters, genotype by location within clusters interaction, year by clusters interaction, genotype by cluster interaction, locations nested in clusters by year interaction, and genotype by year by location within clusters, respectively. **X***_r_*, and **Z***_lr_* up to **Z***_g.y.lr_*, are incidence matrices for their respective effects and dimensions. The remaining model terms were previously defined.

Estimates of variance components from Model 4 were used to obtain CR/DR:

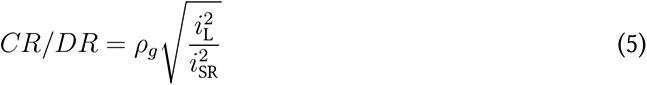

where *ρ_g_* is the correlation between estimated genotypic effects in the non-clustered and clustered sets of locations and 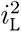 and 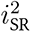 are the estimated reliabilities of genotype means in the non-clustered and clustered sets of locations, respectively. If *CR/DR <* 1, the response to selection will be more effective if selections are made within clusters (Atlin *et al*. 2000a; Bustos-Korts 2017). Note that it is possible for *CR/DR >* 1, indicating that selection will be more effective if the selection is based on eBLUE values obtained from non-clustered locations (Atlin *et al*. 2000a). The terms in equation 5 are defined as follows:

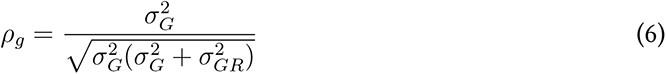

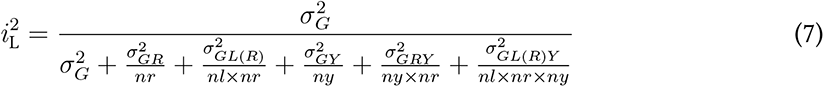

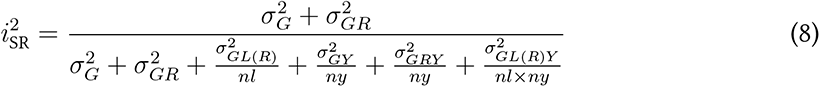

where 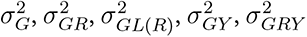, and 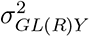 are the genotypic, genotype by cluster, genotype by location nested in cluster, genotype by year, genotype by clusters by year, and genotype by location nested in cluster by year variance components, respectively. *nr* and *ny* are the harmonic means for the number of clusters and years in which genotypes were observed, respectively. *nl* is the median number of locations, obtained from harmonic means within clusters. The residual terms in both reliability estimators were omitted due to the lack of replicated data within environments.

The Jaccard similarity coefficient was used to compare the coincidence of locations within clusters across clustering types. To explore the dynamic for each clustering type, we reported estimates of 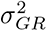 with the jackknife approach presented in section 5.3.4. Lastly, to evaluate if locations were allocated to clusters by chance, we randomly assigned locations into two, three, and four clusters, and assessed their CR/DR. This process was repeated 100 times.

## 6 Results

### 6.1 Single-trial and location analysis

The number of evaluated genotypes by year including checks ranged from 126 to 233, and the number of locations per year ranged from 10 to 32 (Table C3). Estimates of *i*^2^ from single-trial analyses had a median value of 0.55, and the CV% a median value of 7.60% (Figure 1-A). Across years, there has been a positive trend for seed yield with an average increase of 0.49 bu/ac per year (Figure 1-B), although the relative contributions of genetic and non-genetic factors to this trend require further analyses and is the subject of future research.

**Figure 1:**
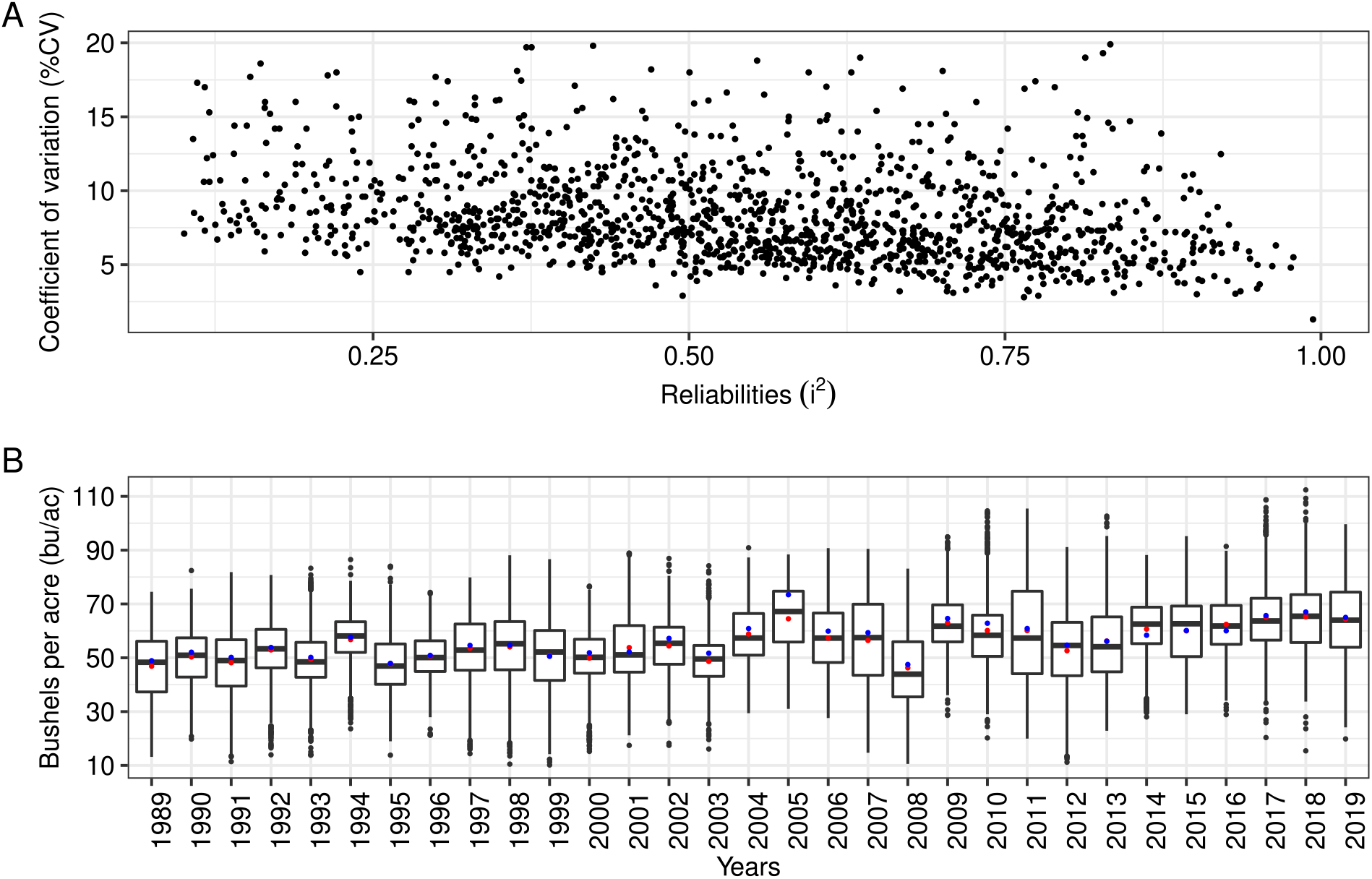
Estimates of reliabilities (*i*^2^) and coefficient of variation (%CV) for 1423 soybean field trials conducted from 1989 to 2019 (A), and boxplots of empirical best linear unbiased estimates (eBLUEs) of seed yield plotted by year (B) from 1989 to 2019. Red dots in B depict the average yield of experimental cultivars excluding checks, whereas blue dots depict the average yield of the check varieties.

### 6.2 Variance components

Most of the estimated phenotypic variance for seed yield has been due to location 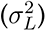 and location by year interaction 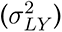 effects (Table 2). Among the estimated GEI variance components 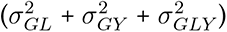, the static contribution 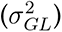 represented 26.30% (Table 2, Model M2-1).

**Table 2:** Point estimates and standard error of variance components for seed yield computed from Soybean Cooperative Tests (1989-2019) using Model 2-1 (baseline) and six clustering methods for clustering locations into mega-environments using Model 4.

| Variance components<sup>a</sup> | Model 2-1 | Clustering criteria<sup>b</sup> |  |  |  |  |  |
| --- | --- | --- | --- | --- | --- | --- | --- |
|  |  | PHE | SoilE | Lat2 | Lat3 | WA | WW |
| $\hat{\sigma}_G^2$ | 6.3 (0.3) | 4.5 (0.4) | 5.0 (0.5) | 4.8 (0.4) | 5.6 (0.4) | 5.7 (0.4) | 6.3 (0.4) |
| $\hat{\sigma}_L^2$ | 58.9 (13.1)<sup>c</sup> | 56.4 (12.6) | 58.6 (13.1) | 59.5 (13.2) | 48.9 (11.4) | 59.1 (13.2) | 60.5 (13.5) |
| $\hat{\sigma}_Y^2$ | 15.3 (5.4) | 13.4 (6.6) | 10.6 (7.2) | 11.5 (6.3) | 15.1 (6.1) | 16.5 (6.5) | 15.0 (6.4) |
| $\hat{\sigma}_{LY}^2$ | 81.4 (5.2) | 74.6 (5.0) | 79.9 (5.2) | 76.5 (5.0) | 77.7 (5.2) | 80.0 (5.1) | 79.9 (5.2) |
| $\hat{\sigma}_{GL}^2$ | 4.8 (0.3) | - | - | - | - | - | - |
| $\hat{\sigma}_{GY}^2$ | 3.2 (0.3) | 2.8 (0.3) | 2.9 (0.4) | 3.2 (0.3) | 3.1 (0.3) | 2.8 (0.3) | 3.0 (0.3) |
| $\hat{\sigma}_{GLY}^2$ | 10.1 (0.3) | - | - | - | - | - | - |
| $\hat{\sigma}_{RY}^2$ | - | 12.2 (5.7) | 7.6 (6.5) | 10.2 (5.2) | 5.9 (3.9) | 2.8 (3.7) | 3.6 (4.1) |
| $\hat{\sigma}_{GR}^2$ | - | 4.1 (0.3) | 2.9 (0.4) | 2.5 (0.3) | 1.5 (0.2) | 1.1 (0.3) | 0.0 (0.0) |
| $\hat{\sigma}_{GL(R)}^2$ | - | 2.5 (0.3) | 4.6 (0.3) | 3.6 (0.3) | 3.8 (0.3) | 4.4 (0.3) | 4.8 (0.3) |
| $\hat{\sigma}_{GRY}^2$ | - | 0.9 (0.2) | 0.5 (0.3) | 0.1 (0.2) | 0.1 (0.2) | 0.7 (0.3) | 0.5 (0.2) |
| $\hat{\sigma}_{GL(R)Y}^2$ | - | 9.7 (0.3) | 9.9 (0.3) | 10.3 (0.3) | 10.2 (0.3) | 9.9 (0.3) | 9.9 (0.3) |
| Clusters | - | 3 | 2 | 2 | 3 | 2 | 2 |

<sup>a</sup> Genotypic ( $\hat{\sigma}_G^2$ ), location ( $\hat{\sigma}_L^2$ ), year ( $\hat{\sigma}_Y^2$ ), location by year ( $\hat{\sigma}_{LY}^2$ ), genotype by location ( $\hat{\sigma}_{GL}^2$ ), genotype by year ( $\hat{\sigma}_{GY}^2$ ), genotype by location by year ( $\hat{\sigma}_{GLY}^2$ ), cluster by year ( $\hat{\sigma}_{RY}^2$ ), genotype by cluster ( $\hat{\sigma}_{GR}^2$ ), genotype by location nested in cluster ( $\hat{\sigma}_{GL(R)}^2$ ), genotype by cluster by year ( $\hat{\sigma}_{GRY}^2$ ), and genotype by location nested in cluster by year ( $\hat{\sigma}_{GL(R)Y}^2$ ).
<sup>b</sup> Phenotypic (PHE), soil and elevation (SoilE), latitude split into two groups (Lat2), latitude split into three groups (Lat3), weather from means across years (WA), and weather from means within years (WW).
<sup>c</sup> Locations were modeled as a random effect in Model 2-1 (Table 1).

Estimated variance components reveal distinct multi-modal distributions over time (Figure 2). For example, the estimated genotypic variance 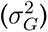 more than doubled from *∼* 3.54 (bu/ac)^2^ for the period 1989-1995 to *∼* 7.56 (bu/ac)^2^ for the period 1996-2003. While the smallest magnitudes of estimated GEI variance components were usually associated with the period from 1989 to 1995, subsequent changes across years were unique to each estimated variance component. A similar pattern was observed for empirical estimates of location, year, and location by year interaction variances (Figure C1).

**Figure 2:**
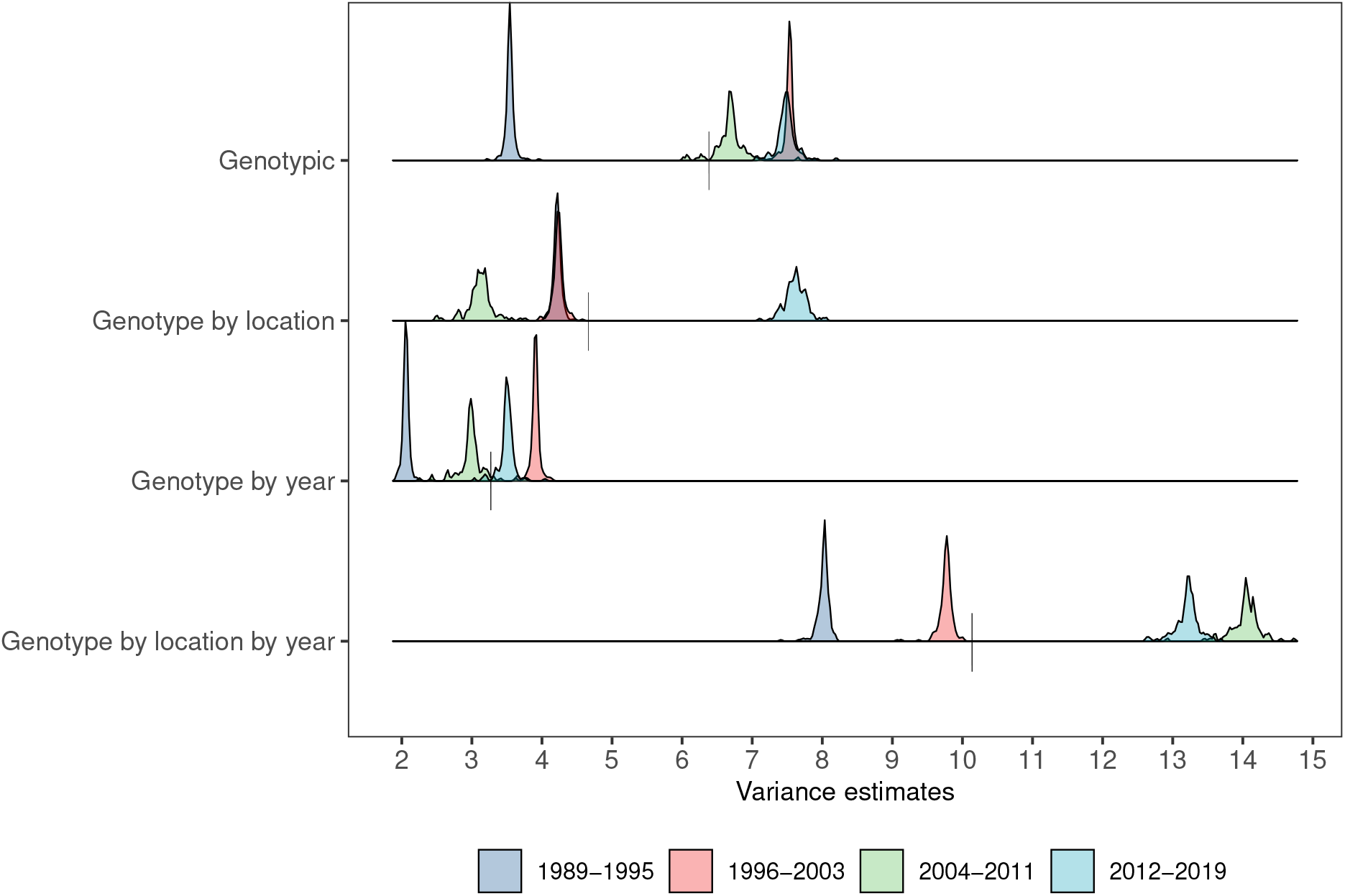
Empirical distributions of estimated variance components consisting of genotypic, genotype by location, genotype by year, and genotype by location by year variances for groups of years 1989-1995, 1996-2003, 2004-2011, and 2012-2019. Empirical estimates were obtained using a jack- knife leave-one-location-out method. Vertical bars on the x-axis represent point estimates across all years.

The best-fit models for the trend of variance components consisted of a mixture of five Log- Logistic distributions for the empirical distribution of 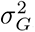, three Log-Logistic distributions for the empirical distribution of 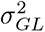, six Log-Logistic distributions for the empirical distribution of 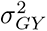, and five Gamma distributions for the empirical distribution of 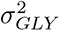 (Table C4, Figure 3). For the empirical distribution of estimated residual variances 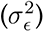, obtained directly rather than through jackknife resampling, the best-fit model was a univariate Log-Logistic distribution (Figure 4). Maximum Likelihood estimates for the distributional model parameters are reported for the selected models (Table 3), as well as plots with empirical and fitted cumulative distribution functions (Figures C2 and C3).

**Figure 3:**
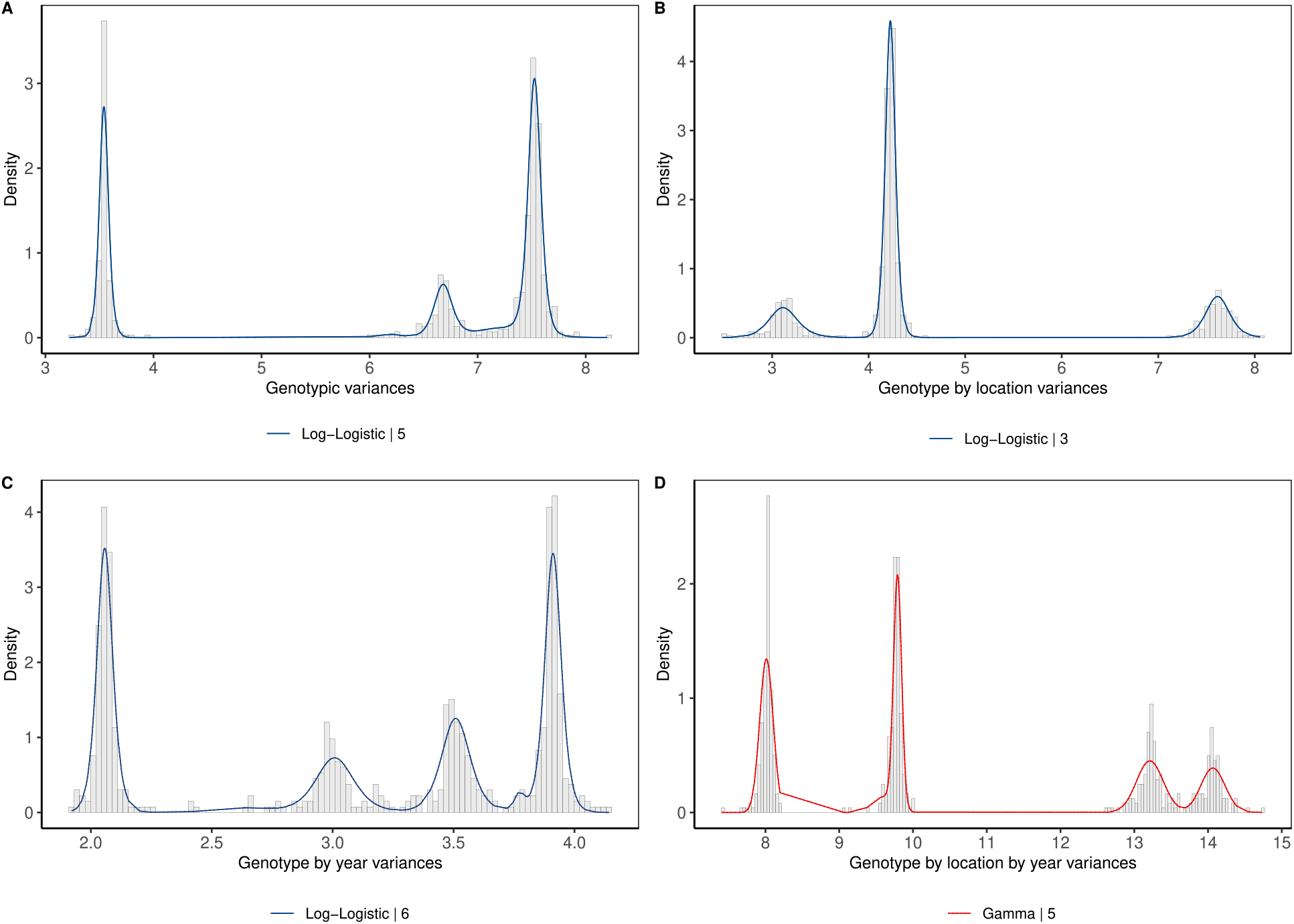
Empirical distributions and probability density function (PDF) of variance components for genotypic (A), genotype by location (B), genotype by year (C), and genotype by location by year (D). Empirical estimates were obtained using a jackknife leave-one-environment-out method. The best-fit PDF models are presented with different colors and include the distribution’s name and its mixture number.

**Figure 4:**
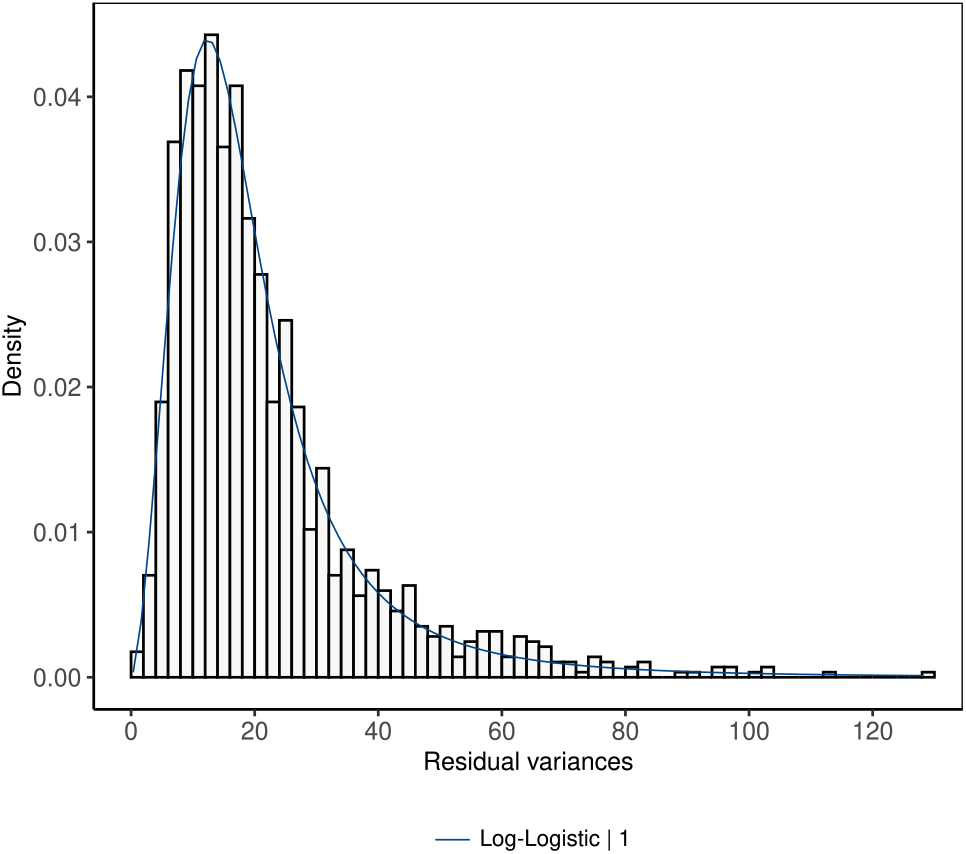
Empirical distribution and probability density function (PDF) for the unimodal Log- Logistic model of residual estimates from individual trials.

**Table 3:** Maximum Likelihood estimates of parameters for the best-fit univariate and multivariate probability distributions for empirical distributions obtained using jackknife resampling. Estimates of residual variance 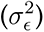 were obtained from trials conducted from 1989 to 2019.

| Variance<sup>a</sup> | Distribution | Number of distributions | Parameters<sup>b</sup> |  |  |
| --- | --- | --- | --- | --- | --- |
| | | | $\hat{w}_i$ | $\hat{\alpha}_i$ | $\hat{\beta}_i$ |
| $\sigma_G^2$ | Log-Logistic | 5 | 0.30 | 125.98 | 3.54 |
|  |  |  | 0.14 | 119.87 | 6.68 |
|  |  |  | 0.08 | 44.03 | 7.23 |
|  |  |  | 0.01 | 88.03 | 6.18 |
|  |  |  | 0.47 | 192.69 | 7.53 |
| $\sigma_{GL}^2$ | Log-Logistic | 3 | 0.63 | 122.27 | 4.23 |
|  |  |  | 0.17 | 32.00 | 3.12 |
|  |  |  | 0.20 | 92.56 | 7.62 |
| $\sigma_{GY}^2$ | Log-Logistic | 6 | 0.02 | 258.41 | 3.77 |
|  |  |  | 0.31 | 172.61 | 3.91 |
|  |  |  | 0.20 | 89.05 | 3.51 |
|  |  |  | 0.30 | 94.97 | 2.06 |
|  |  |  | 0.02 | 32.81 | 2.64 |
|  |  |  | 0.15 | 57.79 | 3.01 |
| $\sigma_{GLY}^2$ | Gamma | 5 | 0.05 | 4078.04 | 0.0024 |
|  |  |  | 0.31 | 7755.00 | 0.0011 |
|  |  |  | 0.20 | 5306.62 | 0.0025 |
|  |  |  | 0.28 | 31922.50 | 0.0003 |
|  |  |  | 0.16 | 7301.02 | 0.0020 |
| $\sigma_\epsilon^2$ | Log-Logistic | 1 | 1 | 2.56 | 17.03 |

<sup>a</sup> Genotypic ( $\sigma_G^2$ ), genotype by location ( $\sigma_{GL}^2$ ), genotype by year ( $\sigma_{GY}^2$ ), and genotype by location by year ( $\sigma_{GLY}^2$ ) variance components.
<sup>b</sup> Estimates of weight parameters ( $w_i$ ) sums to one, and both Gamma and Log-Logistic distributions include a shape ( $\alpha_i$ ) and scale ( $\beta_i$ ) parameter.

### 6.3 Identification of mega-environments

The six clustering strategies revealed that the observed 63 locations could be divided into at least two or at most three ME (Table 4). The PHE clustering presented the lowest CR/DR value, with an optimal number of clusters of three, followed by two clusters for SoilE, WW, and WA. For the PHE clustering type, the best-fit model (M2-18) accounted for heterogeneous variances and pairwise correlations for Σ*_gl_* and Σ*_gy_* with FA(5) and FA(2) matrices, explaining 90.2% and 87.2% of the genetic variances, respectively (Table 1 and Figure C5). For all clustering types, both Silhouette and Elbow criteria indicated similar results (Figure C4). The number of locations within clusters ranged from 7 to 54 (Table 4 and Figure 5), with many clusters having common locations. The greatest Jaccard similarity coefficient was between cluster 1 from PHE and 2 from Lat2 (Table C5).

**Figure 5:**
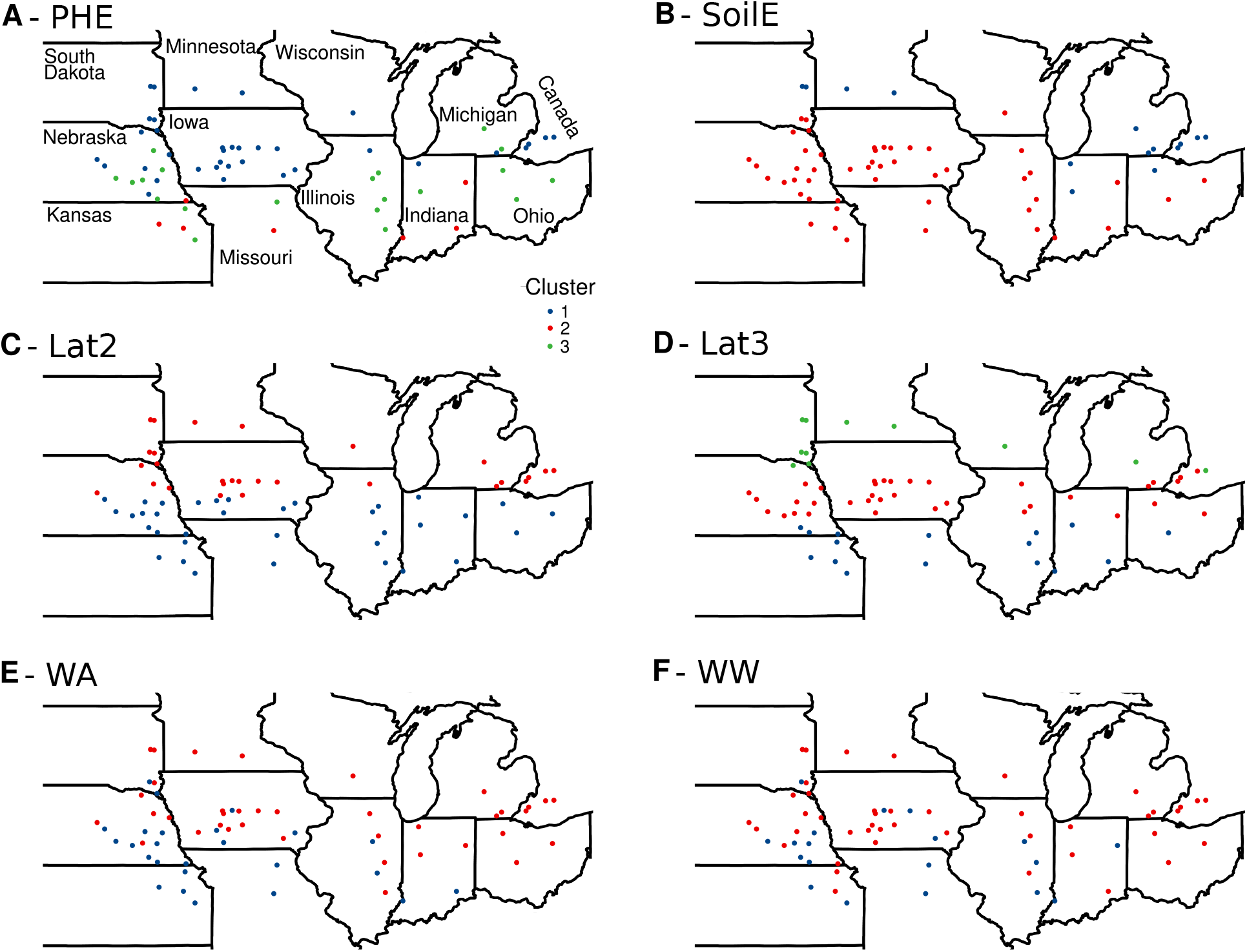
Geographic visualization of the target population of environments divided according to phenotypic (A), soil + elevation (B), latitude split into two groups (C), latitude split into three groups (D), weather across years (E), and weather within years (F) clustering types. In (A), the states’ names are provided for geographic orientation.

**Table 4:** The ratio of correlated responses from selection across all environments relative to direct responses to selection within mega-environments (CR/DR) for each clustering type. *ρ_g_* is the correlation between estimated genotypic effects in the non-clustered and clustered sets of environments, 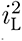 and 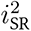 are the reliabilities of genotype means in the non-clustered and clustered sets of environments, respectively.

| Clustering type<sup>a</sup> | Number of |  | Estimates of |  |  |  |
| --- | --- | --- | --- | --- | --- | --- |
| | Clusters | Locations | $\hat{\rho}_g$ | $\hat{i}_L^2$ | $\hat{i}_{SR}^2$ | $\widehat{CR/DR}$ |
| PHE | 3 | 36 / 7 / 20 | 0.72 | 0.38 | 0.51 | 0.62 |
| SoilE | 2 | 9 / 54 | 0.88 | 0.39 | 0.45 | 0.81 |
| Lat2 | 2 | 35 / 28 | 0.81 | 0.39 | 0.48 | 0.73 |
| Lat3 | 3 | 16 / 36 / 11 | 0.89 | 0.44 | 0.44 | 0.89 |
| WA | 2 | 25 / 38 | 0.92 | 0.45 | 0.45 | 0.92 |
| WW | 2 | 19 / 44 | 1.00 | 0.51 | 0.43 | 1.08 |
|  | 2 | - | 0.99 | 0.52 | 0.44 | 1.07 |
| At random | 3 | - | 0.99 | 0.51 | 0.39 | 1.14 |
|  | 4 | - | 0.99 | 0.51 | 0.36 | 1.18 |

<sup>a</sup> Phenotypic (PHE), soil and elevation (SoilE), latitude split into two groups (Lat2), latitude split into three groups (Lat3), weather from means across years (WA), and weather from means within years (WW).

Before K-means clustering, a non-linear PCA was performed with centered and scaled environmental variables (elevation, SV, and MV). The non-linear relationship for most variables was evident from scatterplots (Figures C6, C7, and C8). For SoilE, WA, and WW, the selected number of PC were five (93.06% of variance explained), four (93.42%), and six (91.12%), respectively (results not shown). Furthermore, for MV, just a small proportion of the data was used due to the CERIS biological filter. The selected windows (i.e., with the highest Pearson correlation) were smaller than 20 days, and most of them started at the beginning of the cropping season (Tables C1 and C2, Figure C9).

The effectiveness of clustering was assessed using CR/DR and presented ratios smaller than one in five out of the six cases. These results suggested an improvement in the response to selection when selecting directly within clusters (i.e., regions, ME) versus selecting across all locations for PHE, SoilE, and Lat2, while a moderate response to selection was suggested for Lat3 and WA. For WW, no improvements were predicted. The PHE presented the lowest *ρ_g_* value with an increase in the reliability (from 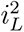 to 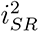, followed by Lat2 and SoilE. For Lat3 and WA, although effective, the reliabilities remained constant. When locations were randomly assigned to two, three, or four clusters, the mean CR/DR values were always bigger than one, with large decreases in the reliability (Table 4).

REML estimates of variances decreased for 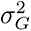 when clusters were included in the complete dataset, except for WW. The observed ratios 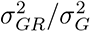 and 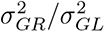 were greather than 0.50 for PHE, SoilE, and Lat2. In terms of the partitioning of 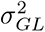, the 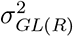 portion was substantially reduced for PHE, Lat2, and Lat3. The 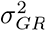 was better captured by PHE, SoilE, and Lat2, being ineffective for WW (estimate bounded at zero due to REML properties). On the other hand, for 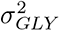, just a small portion of the variation was captured by 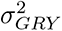 (Table 2). Both PHE and Lat2 clustering types were able to greatly capture 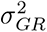 according to analysis for each group of years (Figure C10). Lastly, large reductions in the variation of years 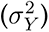 were observed for SoilE and Lat2.

## 7 Discussion

Analyses of balanced data sets produced by MET using the method of moments and REML estimators provide unbiased estimates of variance components. However, data generated by annual PYT and URT are unbalanced and sparse because every year most field plots (experimental units) are reserved for new experimental genotypes, and most previously evaluated experimental genotypes are culled annually. For example, reports of soybean seed yield in MG II and III evaluated in PYT and URT conducted from 1989 and 2019 included less than one percent of all possible combinations of genotypes, locations, and years. Consequently, estimators of variance components might be biased (Rothschild *et al*. 1979). A question to consider is whether the biased estimators produce a large bias in the estimates.

An additional challenge in using soybean seed yield data from PYT and URT is that the data consist of eBLUE values from individual trials within locations for each genotype. These values were transcribed from reports of individual trials formatted as PDF files. To the best of our knowledge, all genotypes were evaluated in replicated field trials organized as RCBD, and were analyzed with a linear model consisting of fixed block and genotypic main effects. To obtain the weights for the stage-wise modeling, the standard errors of genotypic means were assumed to be estimated using equal numbers of replicates per genotype within a field trial. This is equivalent to assuming no missing plots within any field trials, which is highly unlikely. Further, given the lack of replicated data, the estimates of 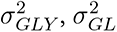 and 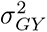 are confounded and biased by 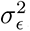, i.e. the plot to plot variability.

A consequence of using eBLUE values instead of individual plot (replicated) data is that estimates of variance components needed to be obtained in multiple stages. If the reports of field trials had provided individual plot data, it would have been possible to produce a variance–covariance matrix associated with adjusted means from each trial. Indeed, if individual plot data are provided, the field plot designs need not be restricted to RCBD wherein the covariances among plots may be substantial and assumptions of independence among plots are inappropriate (MOhring and Piepho 2009). Under such field conditions, the use of spatial models and lattice designs where replicates are considered fixed and blocks as random effects (MOhring *et al*. 2015) can be utilized, and the mixed model framework can provide appropriate weights for analyses of data combined across trials, locations and years. Toward this goal, public soybean breeders are working with curators of “SoyBase” (Drs. Rex Nelson and David Grant, personal communication, 2020) to include results from individual plots in future reports of PYT and URT.

If data were obtained from only URT, they would provide very little information for estimating 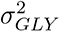 and 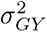 because most experimental genotypes are not grown for more than one year of URT. Although check varieties were replicated across multiple years, the checks are a small sample of genotypes representing mostly commercial cultivars. Thus, inferences about genotypic variability would be limited to the specific sets of checks used in these trials. In addition, by not considering PYT data, the missing data pattern would not be missing at random (MAR). MAR refers to when the missing data pattern in MET is caused by selection (Rothschild *et al*. 1979). If data are not MAR, they can generate biased estimates of variance components (Little and Ru- bin 2020, Chapter 6). Also, URT can have experimental genotypes not included in a previous year of PYT, a condition known as missing completely at random (MCAR) (Little and Rubin 2020; Rubin 1976). Fortunately, recent work from Hartung and Piepho (2021) demonstrated that both MAR and MCAR conditions for field trials conducted in sequential years result in a minor bias for likelihood-based estimators of variance components, a result previously noted for MAR by Piepho and Mohring (2006). Moreover, data from both trials should be included for variance estimation (Laidig *et al*. 2008; Piepho and Mohring 2006), especially to estimate the genotypic variance (Hartung and Piepho 2021).

Despite the caveats mentioned above, the eBLUE values for genotypes from both PYT and URT provided estimates of variance components with similar relative magnitudes as other studies. For example, we determined that environmental, non-genetic sources were the predominant source of variance for seed yield. Similar results were found for wheat in the USA (George and Lundy 2019), winter wheat field trials in Germany between 1983 to 2014 (Laidig *et al*. 2017a), and winter rye (Laidig *et al*. 2017b). Our analyses also revealed that the interactions of genotype with locations 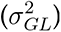 were larger than genotype by year interactions 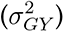. In addition, the three-way interaction among genotype, location, and years 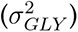, was greater than the two-way interactions involving genotypes. The same pattern was observed when years were combined into four groups of seven to eight years periods. Friesen *et al*. (2016) found similar results for winter wheat evaluated in Canada from 2000 to 2009, where the reported 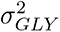 represented 4.1% of the total variation, and both 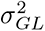 and 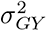 together represented less than 2%. Similar trends were also observed for yield in wheat (George and Lundy 2019; Laidig *et al*. 2017a; Arief *et al*. 2015), rye (Laidig *et al*. 2017b), barley, maize, and sunflower (Laidig *et al*. 2008). While seed yields for multiple crops indicate 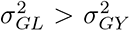, it is not consistent for all traits. For example, Laidig *et al*. (2017b) evaluated the variation in crude protein content, amylogram viscosity, and temperature in winter rye varieties, and reported that 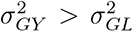, and that the year-to-year 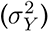 variation was more important than the variation from location to location 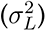. Authors attributed this to the rye seed’s susceptibility towards wetness, low temperature, and radiation during harvest time.

To interpret variance components involving genotypes, we recognize that PYT and URT include genotypes from multiple breeding programs. Each breeding program operates independently with distinct breeding objectives and strategies for markets. For example, a couple of breeding programs have objectives that include high seed yields and greater seed protein for food markets. Seed protein is negatively correlated with seed yield. Thus, the estimates of variance components involving genotypes from these trials are likely greater than for any individual breeding program.

Estimates of variance components were also used to fit parametric probability distributions, in which the data was divided into subsets of seven to eight years. Aguate *et al*. (2019) simulated different MET with variable numbers of genotypes, locations, and years to mimic wheat trials. They found that adding years is more beneficial than adding genotypes or locations for obtaining unbiased estimates of genotypic-related variance components. Furthermore, even in highly imbalanced datasets, estimates from at least eight years of trials produced less than 5% bias in the estimates, compared to biases of 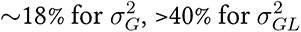, and 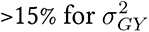, when only two years of MET were considered. Simulation results from Hartung and Piepho (2021) further demonstrated that non-significant bias could be obtained for estimates of 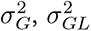, and 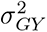, with decreasing dropout rate and increasing the number of years of testing. Both references showed that variance estimates from REML are biased if at least one variance component estimate is bounded at zero, which rarely occurs when using seven or eight years of data. The reader should keep in mind that the goal of minimizing bias in estimators is distinct from the magnitude and proportion of estimates of variance components. The former is a concern for algorithmic estimators while the latter is of concern for breeding decisions about whether to use more locations or more years in their crop species and for traits under selection.

In terms of the fitted probability distributions, especially due to the well-known properties of the analysis of variance (ANOVA) among the breeding community, there is a misconception regarding the distribution of estimates of variance components. If the underlying population is normally distributed, the mean squares are distributed as a chi-square (*χ*^2^), whereas normality and independence are requirements to compute valid F tests from ANOVA (Rencher and Schaalje 2007, Chapter 5). The *χ*^2^ distribution is further used for inferences about variance uncertainty, but as mentioned, it does assume that the random variable is normally distributed. The computed empirical distributions from jackknife can be of any form (distribution), likewise, other approaches such as hierarchical Bayesian models can be employed to obtain posterior distributions of variance estimates. It should be emphasized, however, that the obtained empirical density functions might not necessarily be the data-generating functions. When testing the fit of distributions, the systematic dropping of the data for curation and cleaning (6.8% of the data points were removed) was not considered.

The cluster analyses showed that the observed locations could be effectively divided into three clusters using FA models. The obtained clusters can be considered disjoint subsets of environments with minimal genotypic crossover interaction (BurgueNo *et al*. 2008; Cooper and DeLacy 1994). BurgueNo *et al*. (2008) analyzed a maize MET dataset from CIMMYT and identified five clusters of environments by fitting a FA(2) model. The authors computed the Euclidean distance between pairs of environments from the estimated loading matrix (**Λ**, rotated with singular value decomposition), and environments were clustered based on complete linkage clustering strategy. With a FA(1) model, Bustos-Korts (2017) analyzed data embracing TPE in Denmark, Germany, The Netherlands, and the United Kingdom. The results suggested improvements in response to selection mostly for Denmark, where the CR/DR ratio was 0.93. More recently, Smith *et al*. (2021) proposed a new way to define groups of environments that exhibit minimal crossover interaction based on FA models. The idea is to take advantage of the traditional interpretation of factor and principal component analysis and classify environments into clusters based on the sign (positive or negative) of the estimated and rotated factor loadings.

The identification of homogeneous groups of locations was also accomplished by considering soil plus elevation (SoilE) and meteorological variables when CERIS was applied across environments (WA). The rationale is that a portion of the GEI results from static, repeatable variation (Crespo-Herrera *et al*. 2021; Yan 2016). It is well-known that temperature is a key driving force in the rate of seasonal plant growth (Setiyono *et al*. 2007), which is why GDD is commonly a base unit in crop models (Holzworth *et al*. 2014). Photoperiod plays a significant role in soybean plant development, notably the change from vegetative to reproductive growth. Floral induction is essentially day length and temperature-independent (i.e. conversion of shoot apical and nodal meristems from a vegetative to floral mode). In soybeans, this induction occurs as soon as the first unifoliolate leaflets emerge and expand, becoming capable of measuring the night length (from dusk to dawn). Once floral induction occurs at a given apical or axillary node, the few-celled vegetative apical zone is transformed from a vegetative development pathway into a floral inflorescence development pathway. The development pathway is back under thermal control (Setiyono *et al*. 2007). Soybean is a quantitative long-night length sensing (not a short-day length sensing), and hence highly influenced by photoperiod and therefore by the latitude of the growing region/trial (Jackson 2009). This is a major reason that different soybean maturity groups are grown at different latitudes (Mourtzinis and Conley 2017). The estimated clusters (except for the *ad-hoc* Lat2 and Lat3) follow a certain pattern in terms of latitude, which was also confirmed by the Jaccard similarity.

For this work, we focused on identifying ME through reliable estimates of variance components. However, other environmental subsets would have been formed using different clustering strategies (BurgueNo *et al*. 2008). Given that our approach is unsupervised learning (i.e. we do not know the truth about ME), the objective is always to discover an interpretable grouping of members. We addressed interpretation using the effectiveness of clustering (Atlin *et al*. 2000a). Other strategies for clustering can also be tested, for example, empirical knowledge of the TPE. But regardless of the definition/identification of ME, breeders can take advantage of the best linear unbiased prediction (BLUP) that borrows information (strength) between ME from the genotype by ME interaction. This type of modeling can be highly beneficial for ME that rely on a small number of locations (Buntaran *et al*. 2021; Piepho *et al*. 2016; Piepho and MOhring 2005).

A natural continuation for this work would be to (i) evaluate the effectiveness of a combined cluster from soil, elevation, and meteorological variables filtered by CERIS; (ii) evaluate if BLUP based-models will improve the selection response upon regionalization in the Uniform Soybean Cooperative Tests; (iii) select an appropriate model that might account for heterogeneous covariances among ME as well genetic relationships, because we confined attention to the compound symmetry model to facilitate comparison of the different clustering types; and (iv) leverage how far back in the historical data should we go to take maximum advantage of the data in current models. It is also worth investigating if modeling maturity groups (especially when more data is considered) would enhance the ability to find meaningful ME using phenotypic models.

## 8 Conclusion

We dissected the sources of soybean seed yield variation using reports from Soybean Cooperative Tests for maturity groups II and III. We determined that the sampled locations can be split into mega-environments according to phenotypic, geographic, and meteorological information. Furthermore, it was possible to monitor trends in variance components involving genotypes in terms of parametric probability distributions. Historical field trials also evaluate traits like seed quality and size, iron deficiency chlorosis, green stem, seed oil, and protein content. Therefore, the approach presented herein can be applied to multiple economically important quantitative traits. It is important to emphasize that our inference space is restricted to germplasm adapted to MG II and III and evaluated as nearly homozygous genotypes in public MET. Also, the evaluated environments represent a random sample of the target population of environments. Thus, inferences about the identified mega-environments (ME) are likewise restricted to the field trial network explored by public soybean breeding programs. Finally, in addition to the practical and theoretical results applied to soybean genetic improvement, the analyses performed in this study may be applied to quantitative traits evaluated in any crop using MET.

## 9 Acknowledgments

Our sincere thanks to Dr. Aaron Lorenz’s group for providing a large portion of the phenotypic data (in particular to Ben Campbell, Liana Nice, and Cleiton Wartha); Dr. Rex Nelson for providing individual plot data from trials conducted in years 2018 and 2019; Dr. Jim Specht for valuable insights on environmental factors that influence soybean growth and development; Dr. Alencar Xavier for careful reading and suggestions on the early draft of the manuscript; and the Iowa State University Research IT team for providing efficient computational power. Most importantly the authors appreciate the long-term commitments of all collaborators (breeders, staff, students, farmers, etc.) that have worked on conducting the cooperative Soybean trials every year since 1983. Funding for this research was provided by the Department of Agronomy - Iowa State University, the North Central Soybean Research Program, an NSF grant (1830478), Baker Center for Plant Breeding, and USDA-ARS CRIS Project IOW04714.

## 11 Author contributions

MDK and WDB conceived the research; MDK performed the statistical analyses and wrote the first drafts of the manuscript; KOGD provided insights into the methodology and helped in the interpretation of the results; AKS provided interpretation of the results and guidance in scientific writing; and WDB and AKS were responsible for acquiring funding to support the research. All authors critically revised drafts of the manuscript and approved the final version.

## 12 Data and code availability

The developed R codes to reproduce most analyses are publicly available on GitHub (https://github.com/mdkrause/VarComp-ME). The data sets (phenotypic, weather, and soil data) are available in the developed R package SoyURT (https://github.com/mdkrause/SoyURT). Raw data can be further downloaded from Soybase (https://soybase.org/ncsrp/queryportal/), and directly from the PDF files from the USDA-ARS webpage (https://www.ars.usda.gov/midwest-area/west-lafayette-in/crop-production-and-pest-control-research/docs/uniform-soybean-tests-northern-region/). All cited web pages here and throughout the manuscript were accessed on 05/04/2023.

## 14 Appendix A - The structure of the data

The complete data consist of eBLUE values for seed yield (bu/ac) of experimental genotypes and check varieties grown in field trials of soybean maturity groups (MG) 00 through IV from 1941 to 2020. For our purposes, we restricted our analyses to data from the most reliable trials of MG II and III from 1989 to 2019. We restricted our interest to MG II and III because these are North America’s primary soybean production maturity groups.

We defined unreliable trials as individual trials with estimates of reliability (*i*^2^) less than 0.10 and coefficients of variation (CV%) greater than 20%. Reliability values less than 0.10 and CV values greater than 20% are indicators of multiple experimental units within a replicate that have been severely damaged by unknown sources (such as deer herds, raccoon troops, hail storms, etc.), causing unexpected, inconsistent phenotypic values between replicated genotypes within the trial at an environment. Not only are data from the damaged experimental units suspect, but adjacent undamaged experimental units have unfair advantages from lack of competition. Thus, in keeping with the GIGO (garbage in - garbage out) maxim, plant breeders routinely exclude data from unreliable field trials prior to making decisions about which genotypes should be retained for future trials and breeding purposes. Our exclusion rules are consistent with best practices by public and commercial soybean breeders.

To assure that individual genotypes with unexplained low yields did not affect the variance estimates, 12 individual records within trials with an estimated mean of less than ten bu/ac were removed. In total, only 6.8% of the observed data points were removed. Lastly, locations with less than three years of data were excluded from our analyses. The resulting data comprised 4,257 experimental genotypes evaluated at 63 locations, in 31 years, resulting in 591 location-year combinations (environments) with 39,006 yield values (data points).

## 15 Appendix B - The analysis within versus across MG

We restricted our analyses to data from the most reliable trials (Appendix A) belonging to MG II and III from 1989 to 2019. Our approach was to make inferences across MG, but here we present the analysis within MG. The results (point estimate followed by the standard error) from the analyses across (presented in Table 2) and within (with individual datasets) MG are the following:

| Variance components | Across MG II and II | MG II | MG III |
| --- | --- | --- | --- |
| Genotypic (G) | 6.3 (0.3) | 5.4 (0.4) | 4.43 (0.4) |
| Location (L) | 58.9 (13.1) | 54.2 (16.1) | 48.8 (15.6) |
| Year (Y) | 15.3 (5.4) | 13.7 (5.9) | 27.9 (10.4) |
| LY | 81.4 (5.2) | 85.7 (6.9) | 87.8 (7.9) |
| GL | 4.8 (0.3) | 4.6 (0.4) | 3.8 (0.4) |
| GY | 3.2 (0.3) | 3.2 (0.3) | 2.8 (0.3) |
| GLY | 10.1 (0.3) | 8.3 (0.4) | 9.8 (0.4) |

The interpretation within MG is the same: GL is more important than GY; and the static portion of the GEI (GL + GY + GLY) due to GL accounted for 26%, 28%, and 23%, respectively. An alternative way to test for homogeneous (pooled) versus heterogeneous variances within MG can be implemented with two different versions of Model 2. The model with heterogeneous variances within MG assumes that there is a unique value for each genotypic-related variance. For example, the estimated genotypic variance using data from both groups is 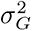, and within MG is 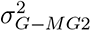 and 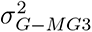, for MG II and MG III, respectively. Both models can be fitted in *Asreml-R* (or any other mixed model package) using the *SoyURT* package as follows:

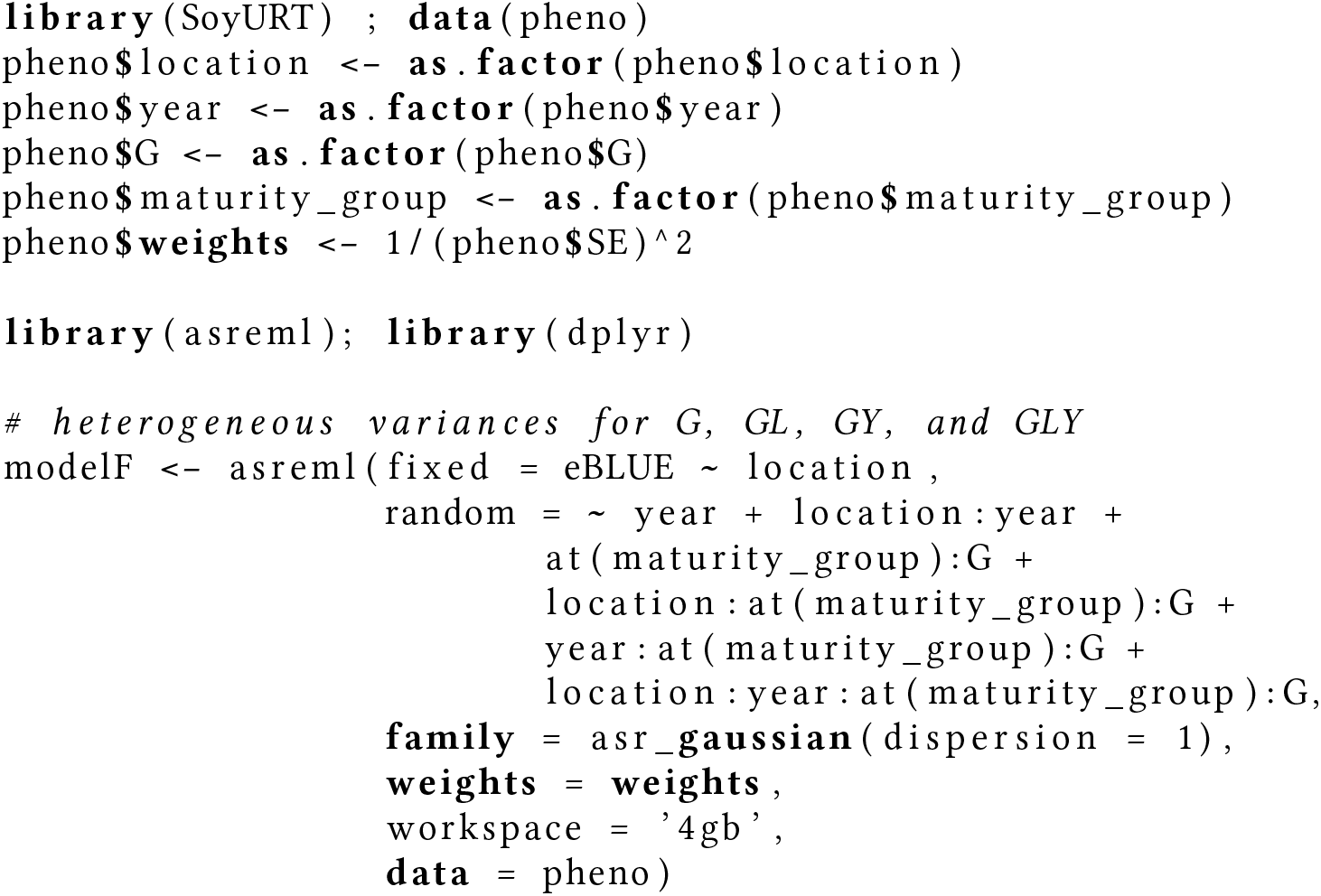
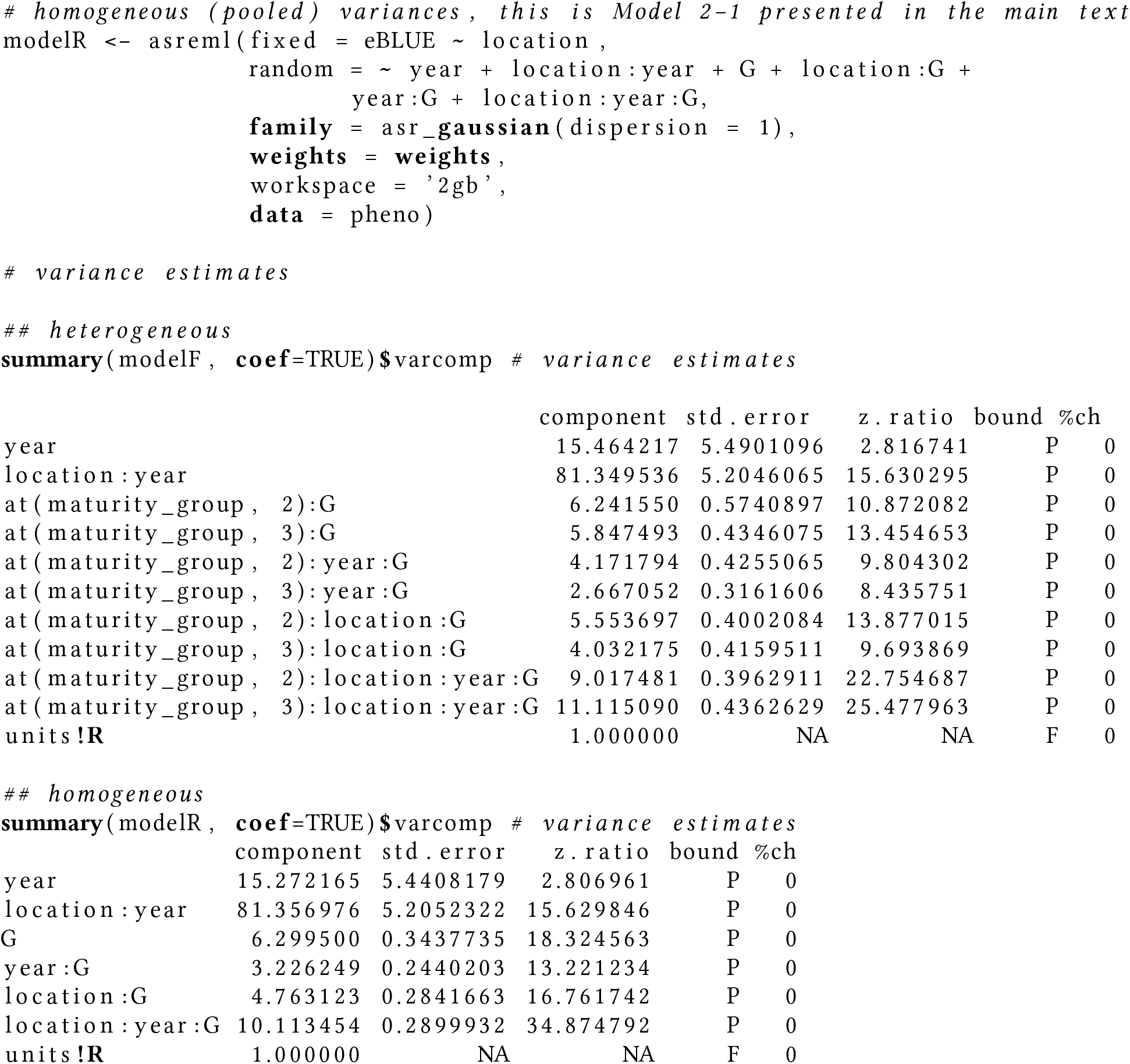

With the fitted models, it is possible to evaluate the AIC/BIC selection criteria to check which model has a better goodness-of-fit:

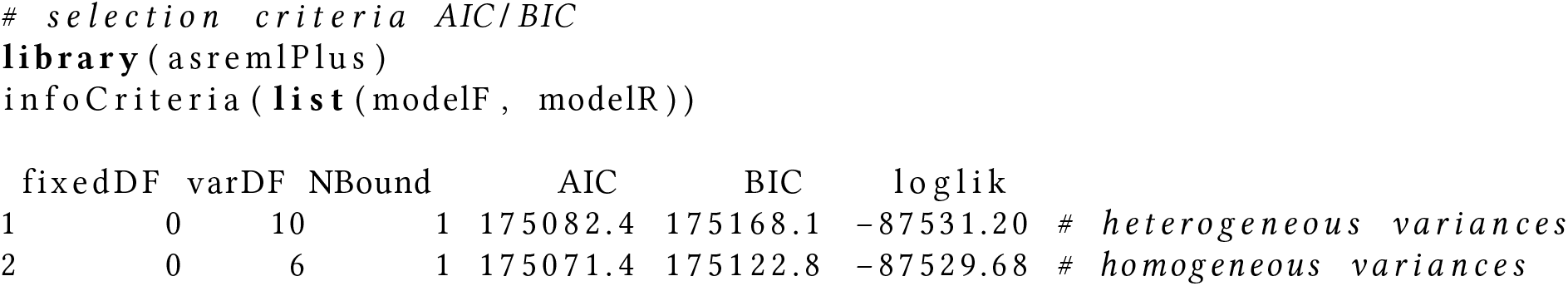

There are ten variance components to be estimated in the heterogeneous model (*modelF*), whereas for the homogeneous model (*modelR*), six variance components. Both AIC and BIC values are presented above. Therefore, there is evidence it is reasonable to use the homogeneous variance model and obtain pooled variance estimates for MG II and III. In addition to the statistical modeling, the challenge, of course, is to delineate a continuous characteristic (maturity) that is affected by the environment (heat and drought). For example, what is the difference between a genotype that has been classified as 2.9 and a genotype that has been classified as 3.1? Is the line classified as 2.9 a member of MG II (consisting of lines classified as 2.1) or better classified as MG III? In other words, experimental lines classified as MG II might actually be early MG III lines, whereas early experimental lines classified as MG III might be late MG II lines. Moreover, by combining both MG, the genetics and the locations were better sampled.

Lastly, we fitted the following model considering MG as a fixed effect:

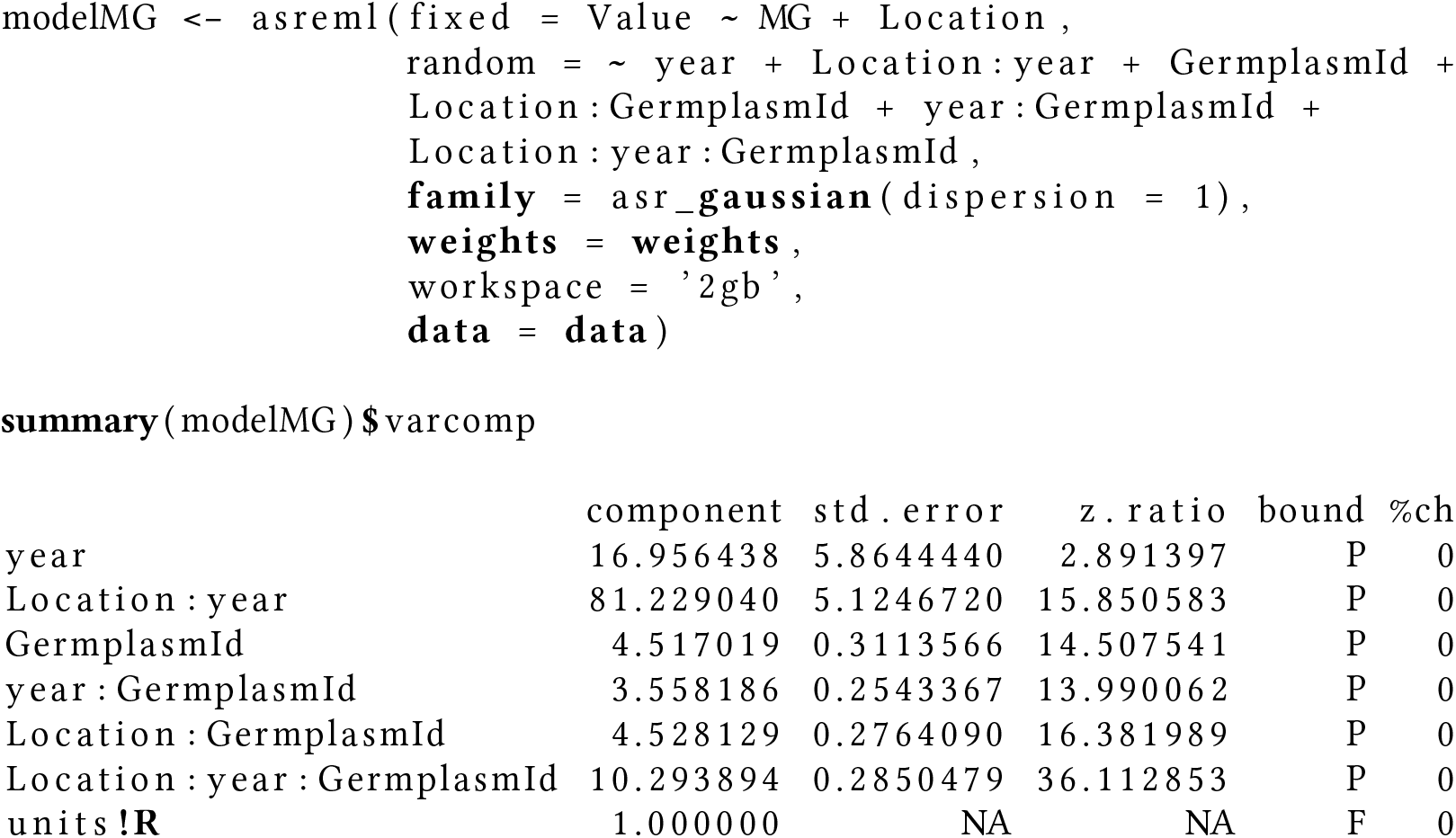

In conclusion, the REML estimates of variance are similar to the ones reported in the main manuscript.

## 16 Appendix C - Additional figures and tables

**Figure C1:**
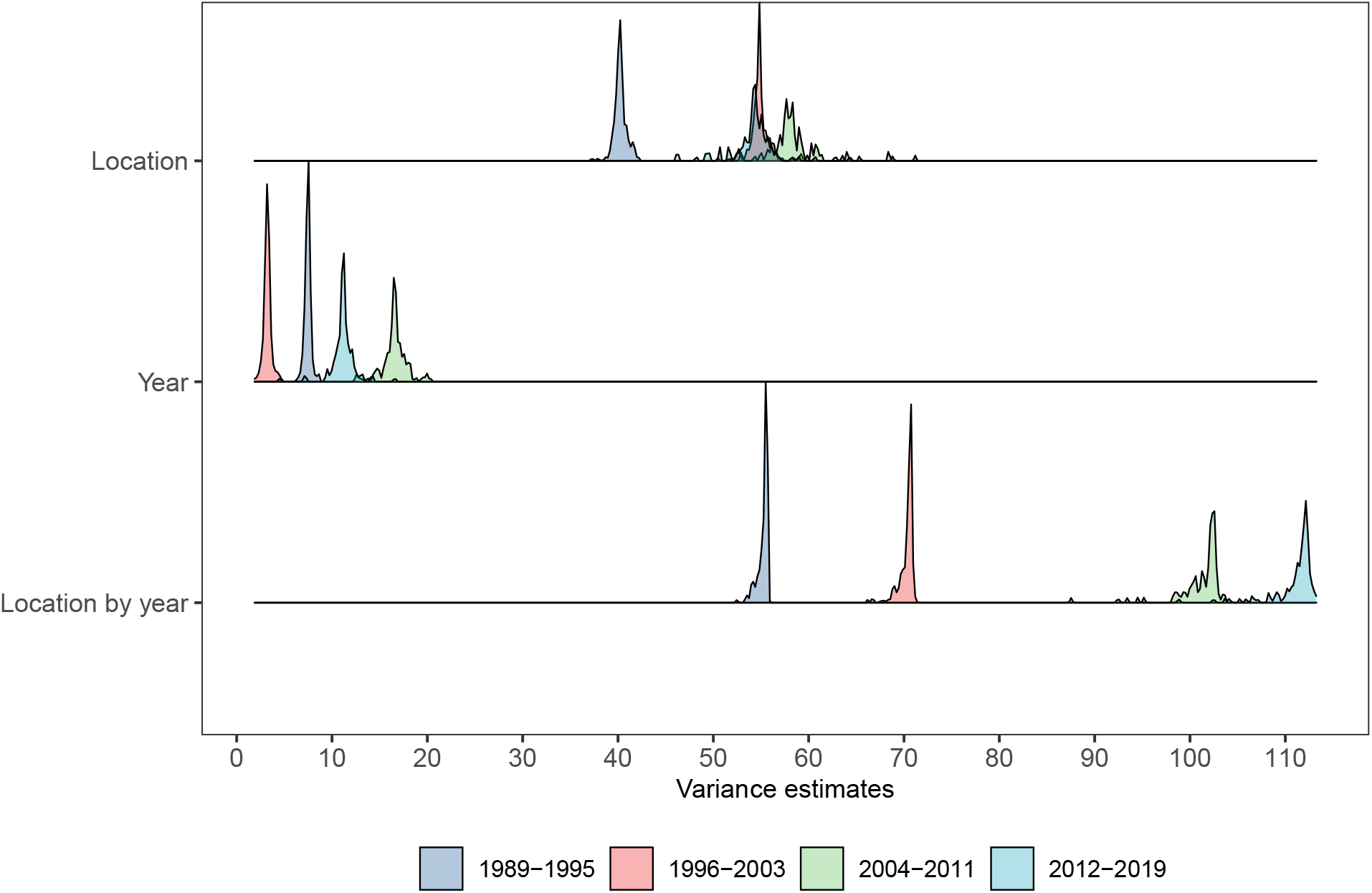
Jackknife estimates of location, year, and location by year variances for the groups of years 1989-1995, 1996-2003, 2004-2011, and 2012-2019.

**Table C1:**
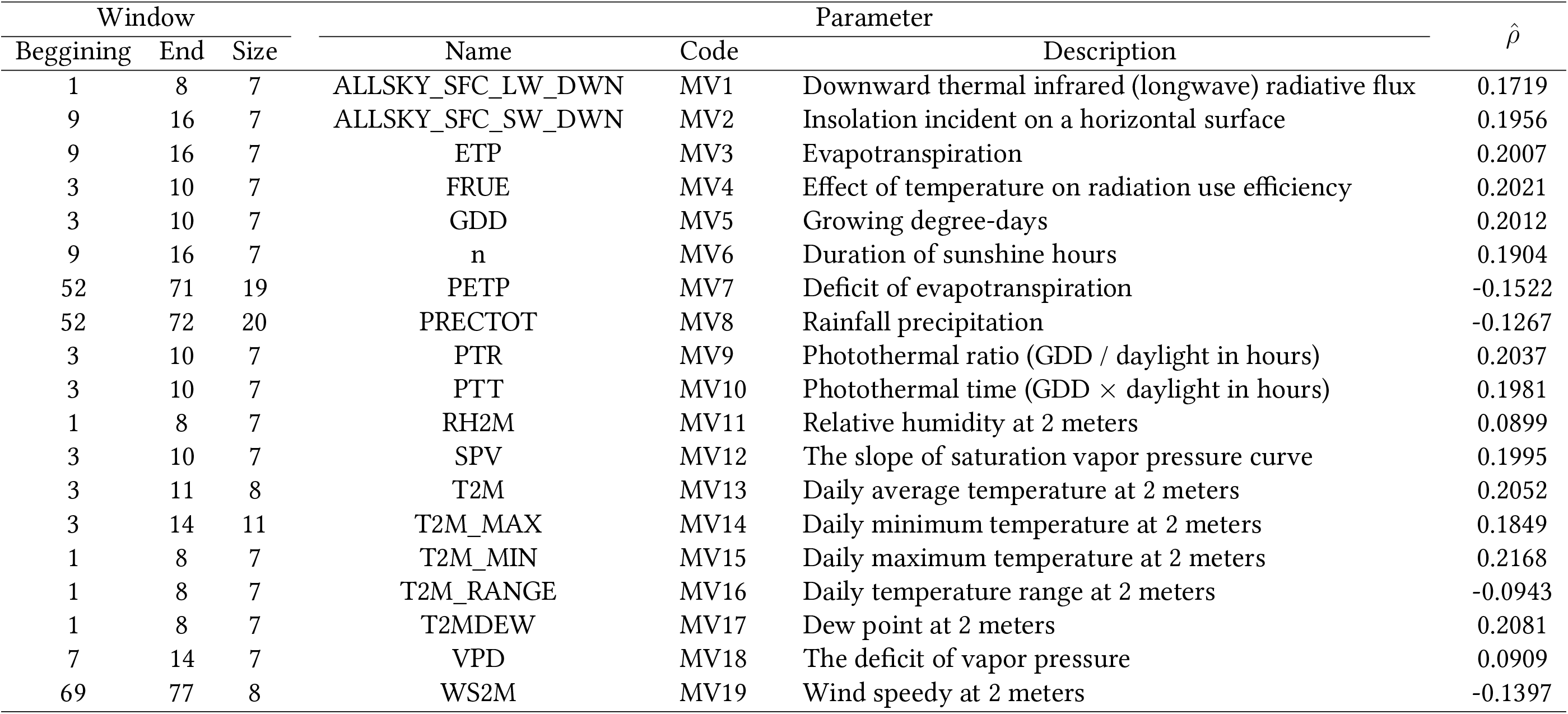
Meteorological variables and selected window across years (WA) with the highest Person correlation (*ρ̂*).

**Table C2:**
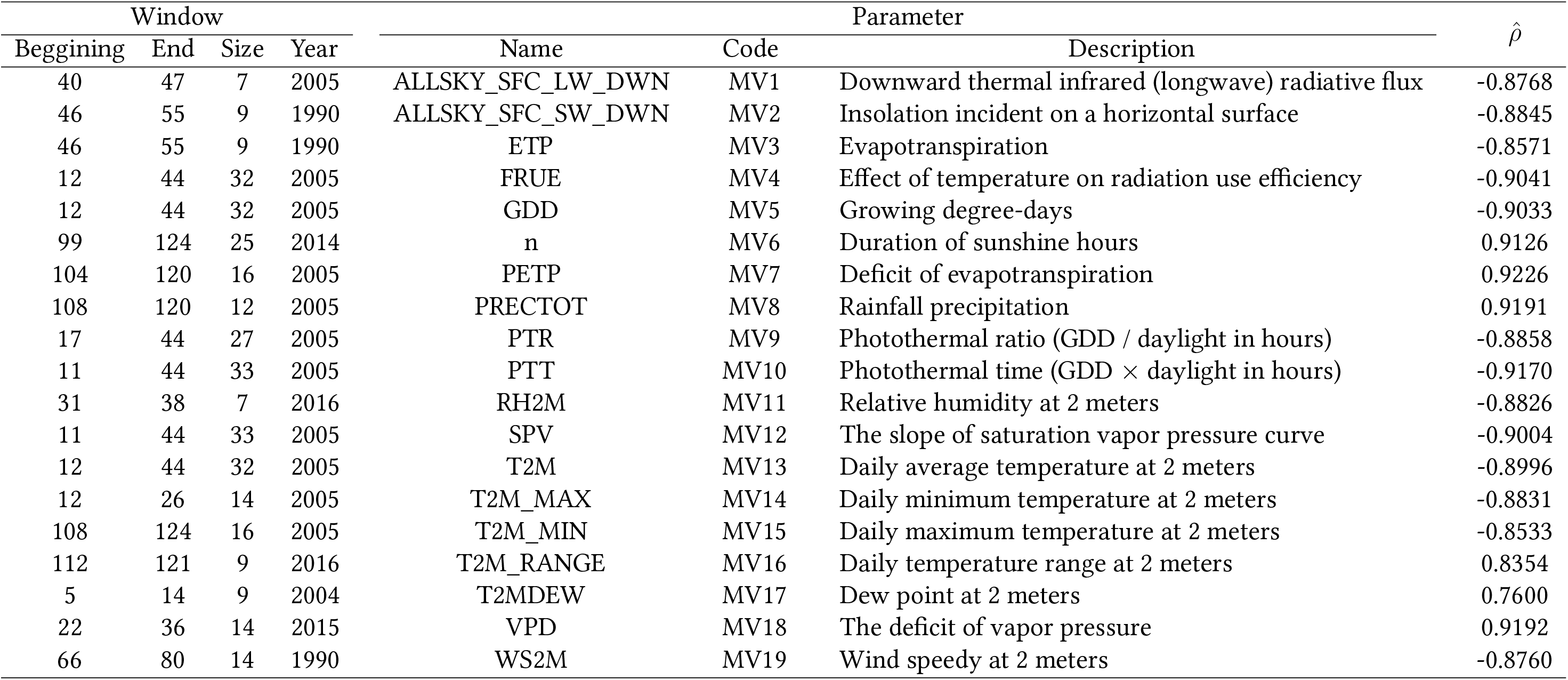
Meteorological variables and selected window within years (WW) with the highest Person correlation (*ρ̂*).

**Figure C2:**
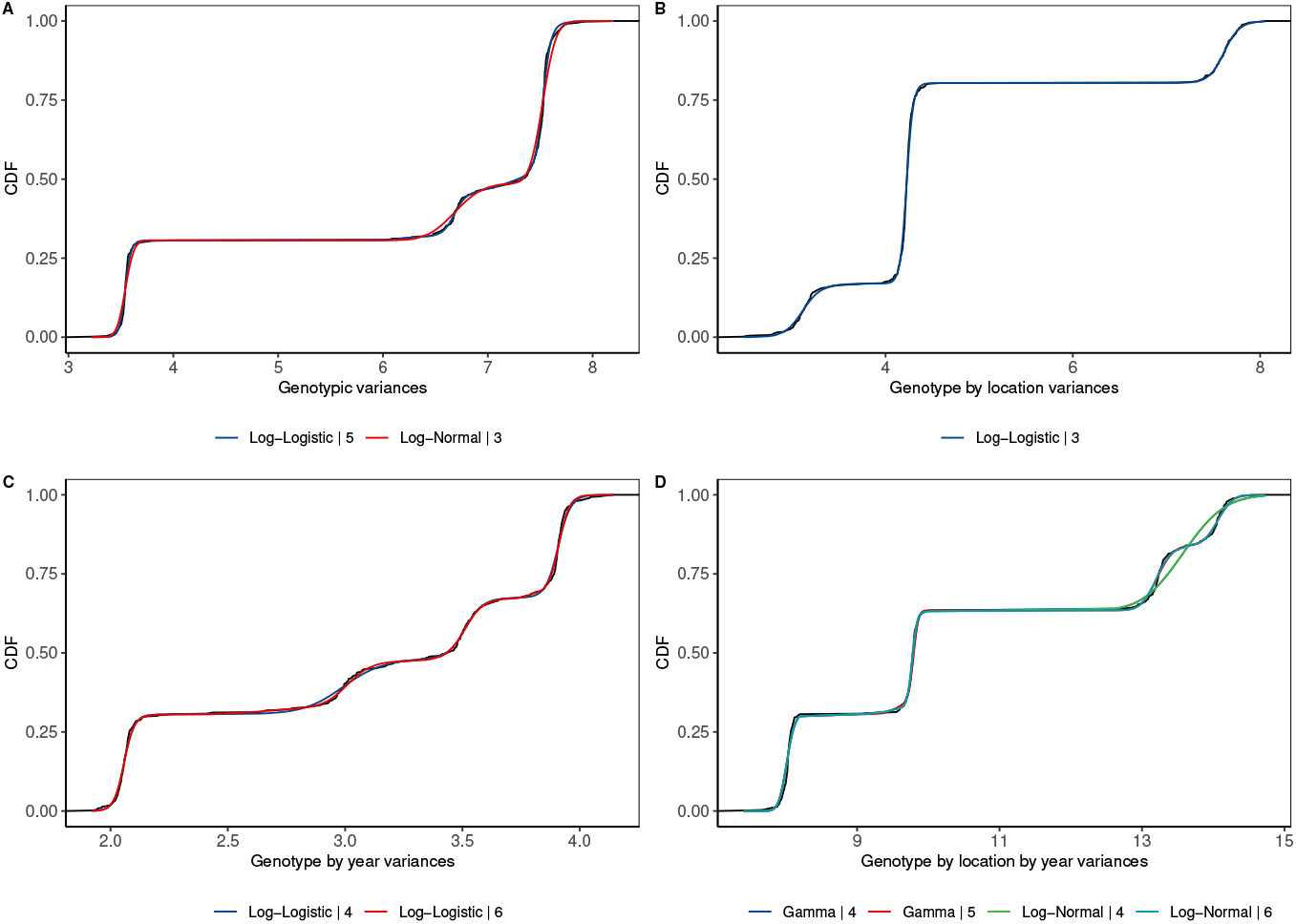
Cumulative distribution function (CDF) for the best-fit models according to the genotypic (A), genotype by location (B), genotype by year (C), and genotype by location by year (D) variances from jackknife. In each plot, the legends states for name of the distribution followed by its mixture number.

**Figure C3:**
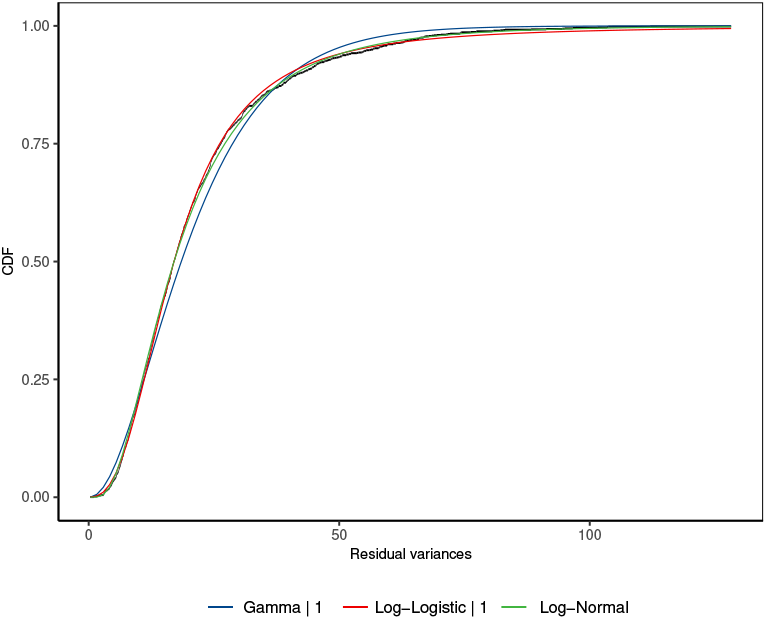
Cumulative distribution function (CDF) for the best-fit models according to the residual variances from individual trial level. The legends state the name of the distribution followed by its mixture number.

**Table C3:**
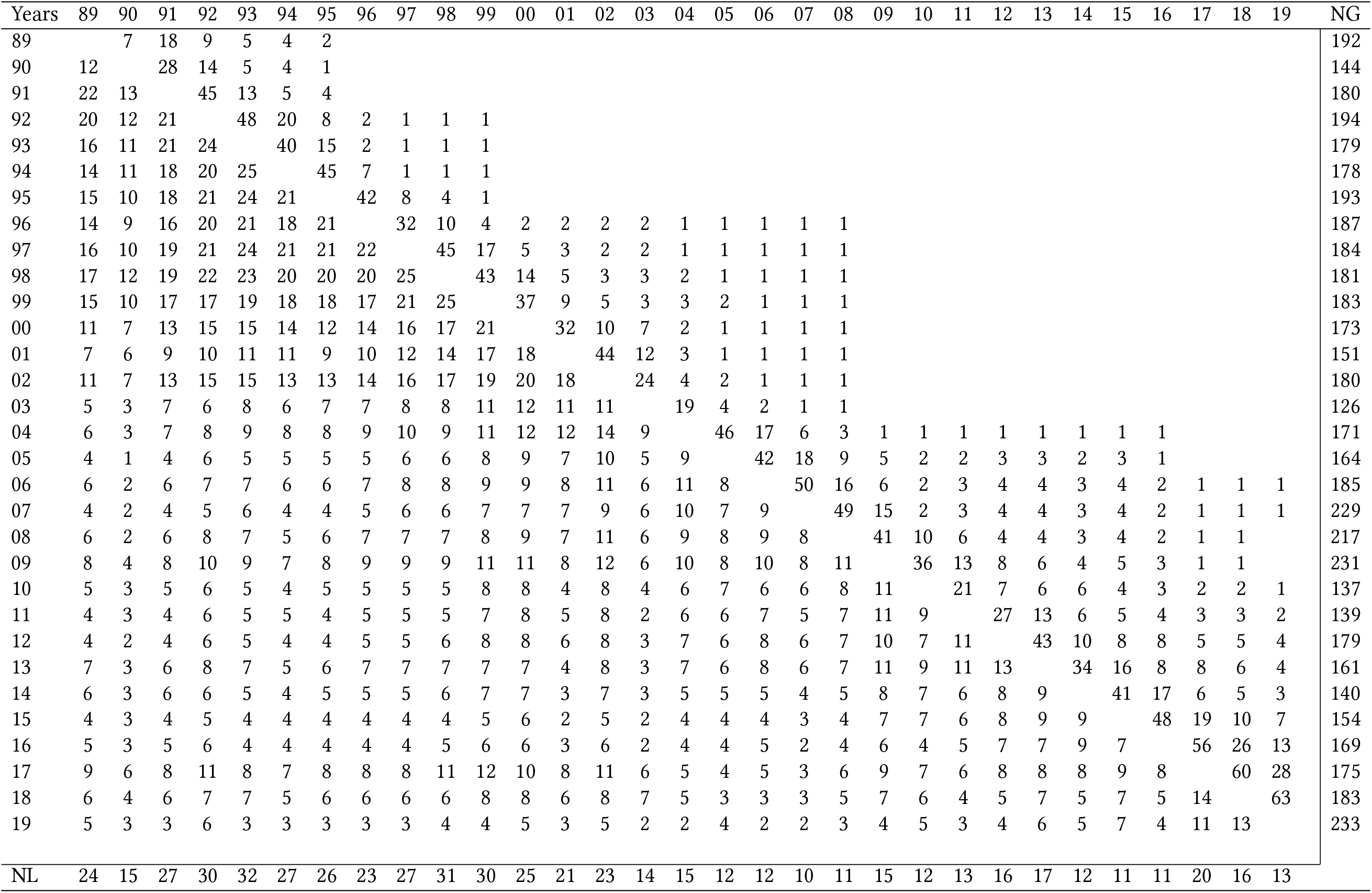
Number of genotypes (NG) and locations (NL) within each year, the number of common genotypes between years (upper diagonal), and the number of common locations between years (lower diagonal).

**Table C4:**
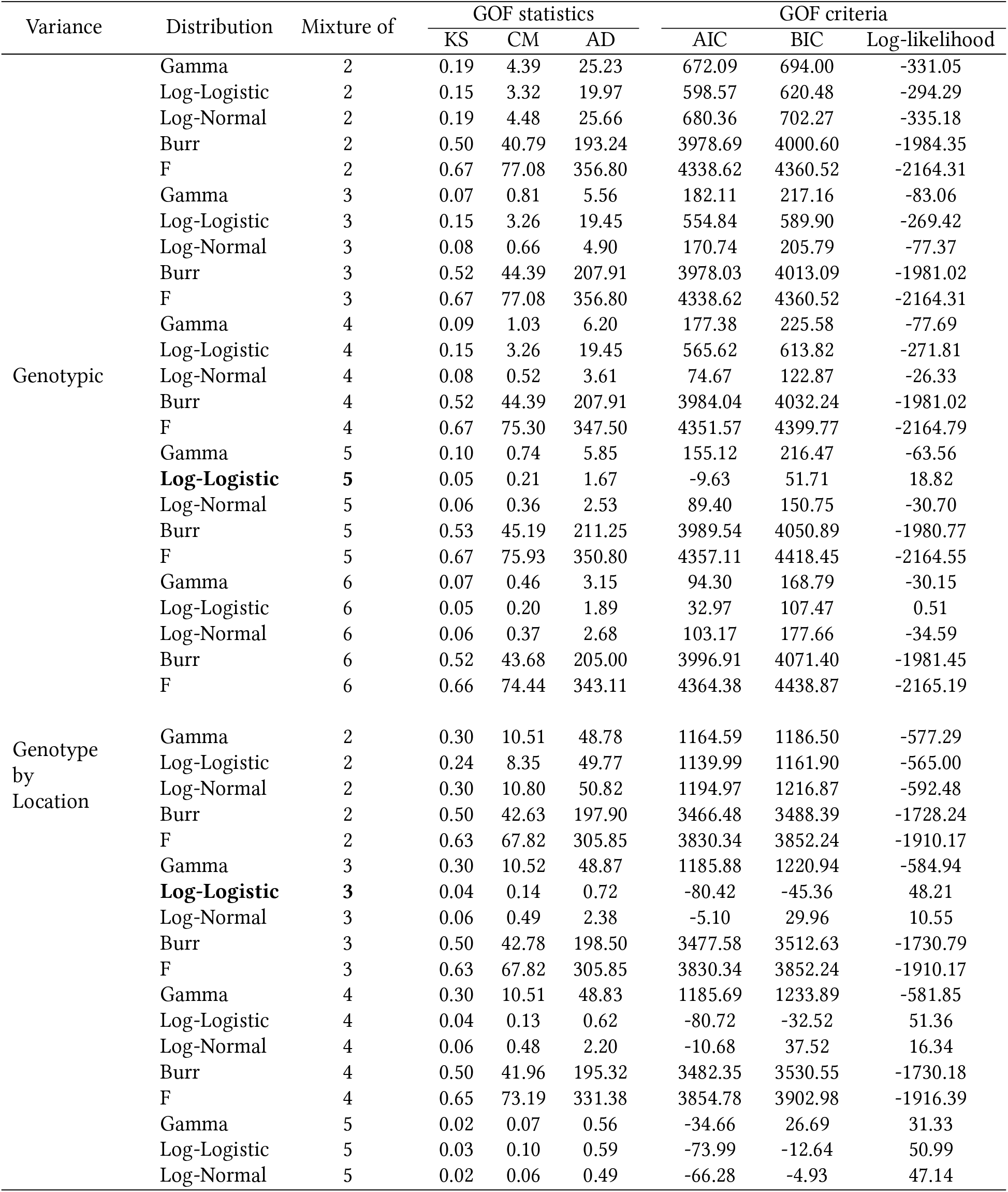
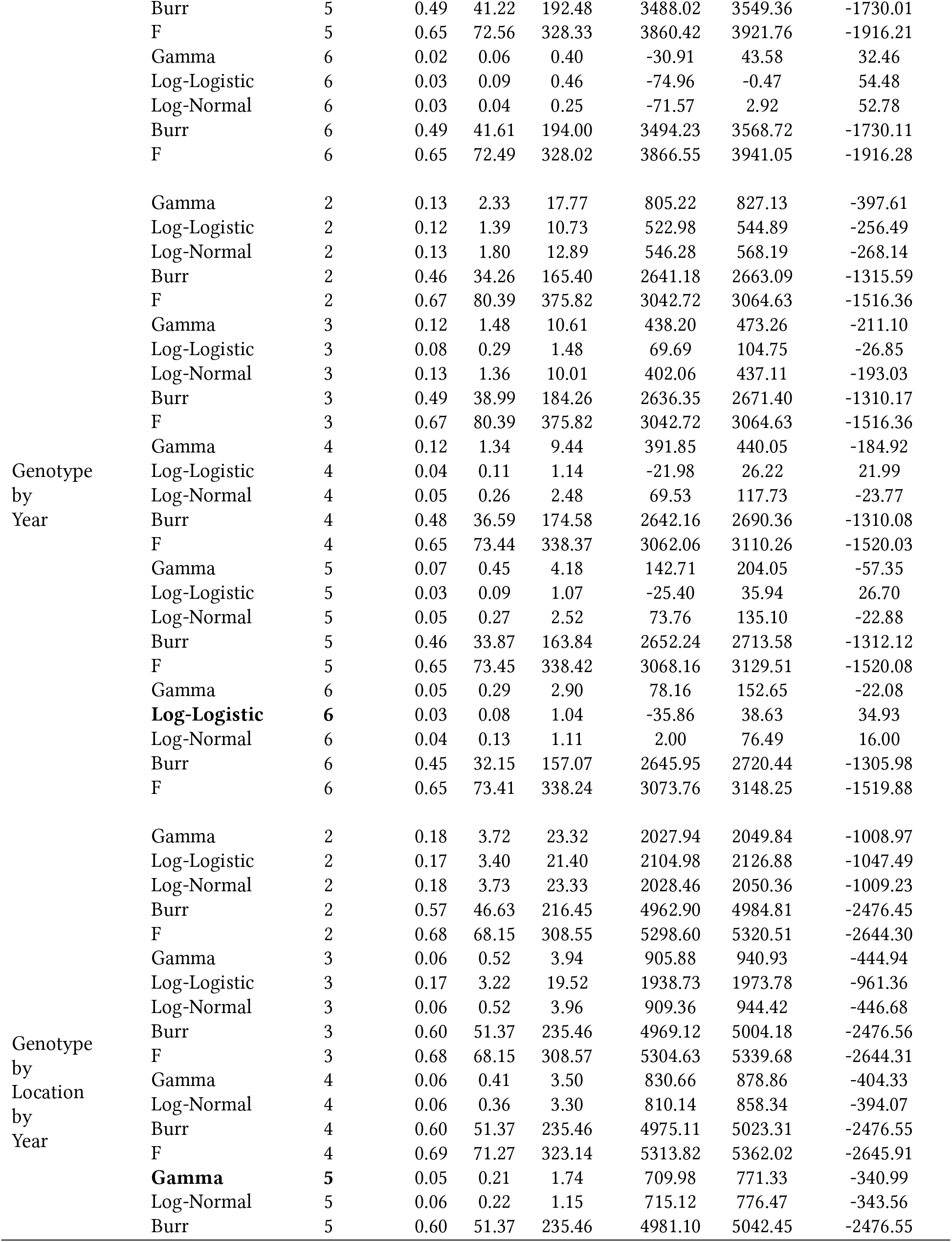
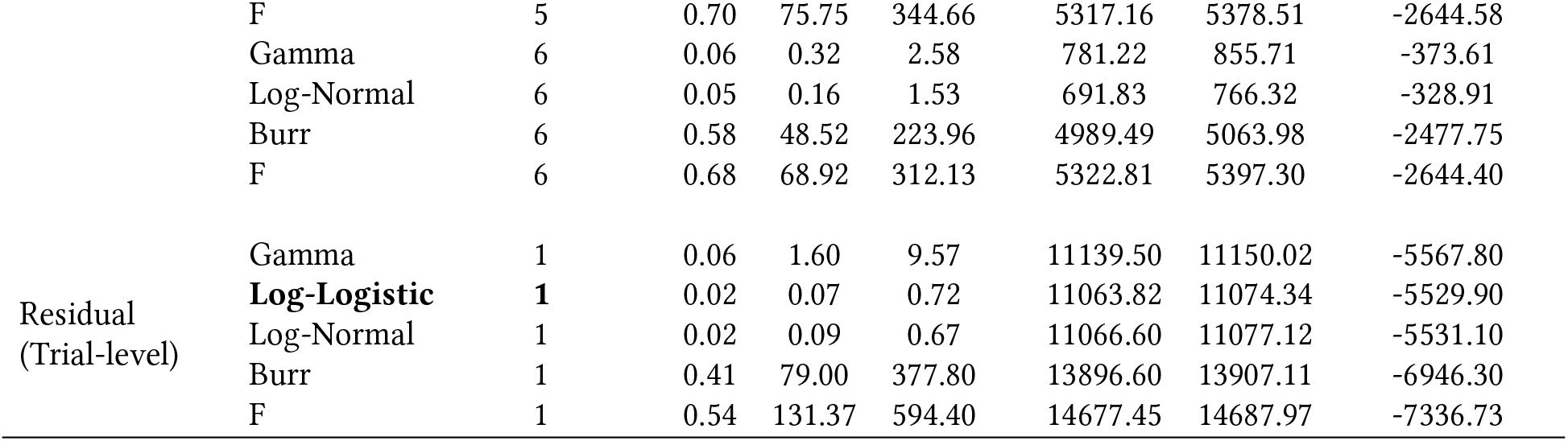
Goodness-of-fit (GOF) statistics and selection criteria for the fit of univariate and multivariate probability distributions for genotypic, genotype by location, genotype by year, and genotype by location by year variance components estimated from the jackknife analysis, and residuals variances from trial-level. The best-fit model is highlighted in bold. KS, CM, AD, AIC, and BIC stand for Kolmogorov-Smirnov, Cramer-von Mises, Anderson-Darling, Akaike’s and Bayesian Information Criterion, respectively.

**Figure C4:**
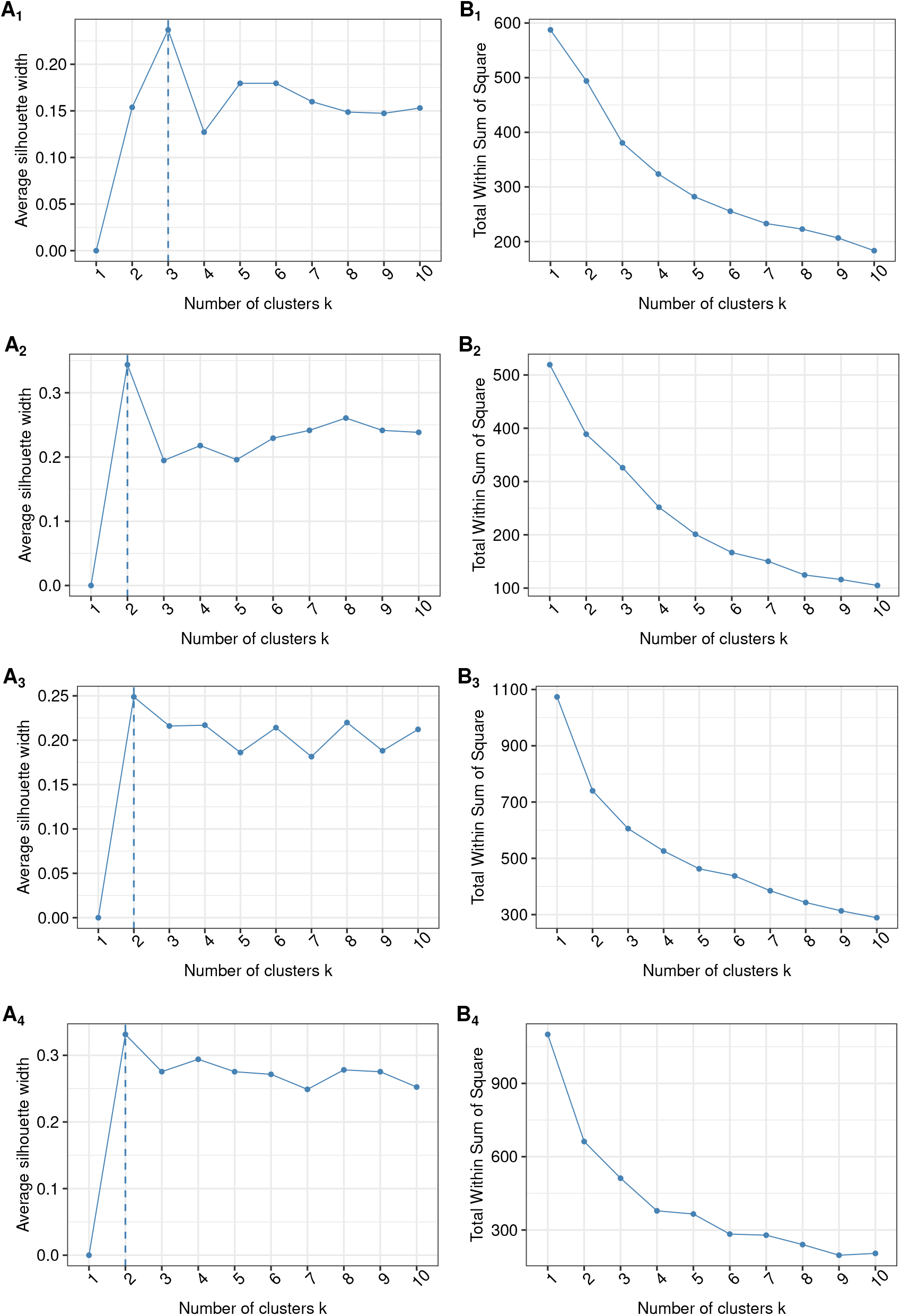
Graphical display of the optimal number of clusters based on the Silhouette (A) and Elbow (B) methods for the phenotypic clustering type (PHE, A_1_ and B_1_), soil + elevation variables (SoilE, A_2_ and B_2_), weather within year variables (WW, A_3_ and B_3_), and weather across year variables (WA, A_4_ and B_4_).

**Figure C5:**
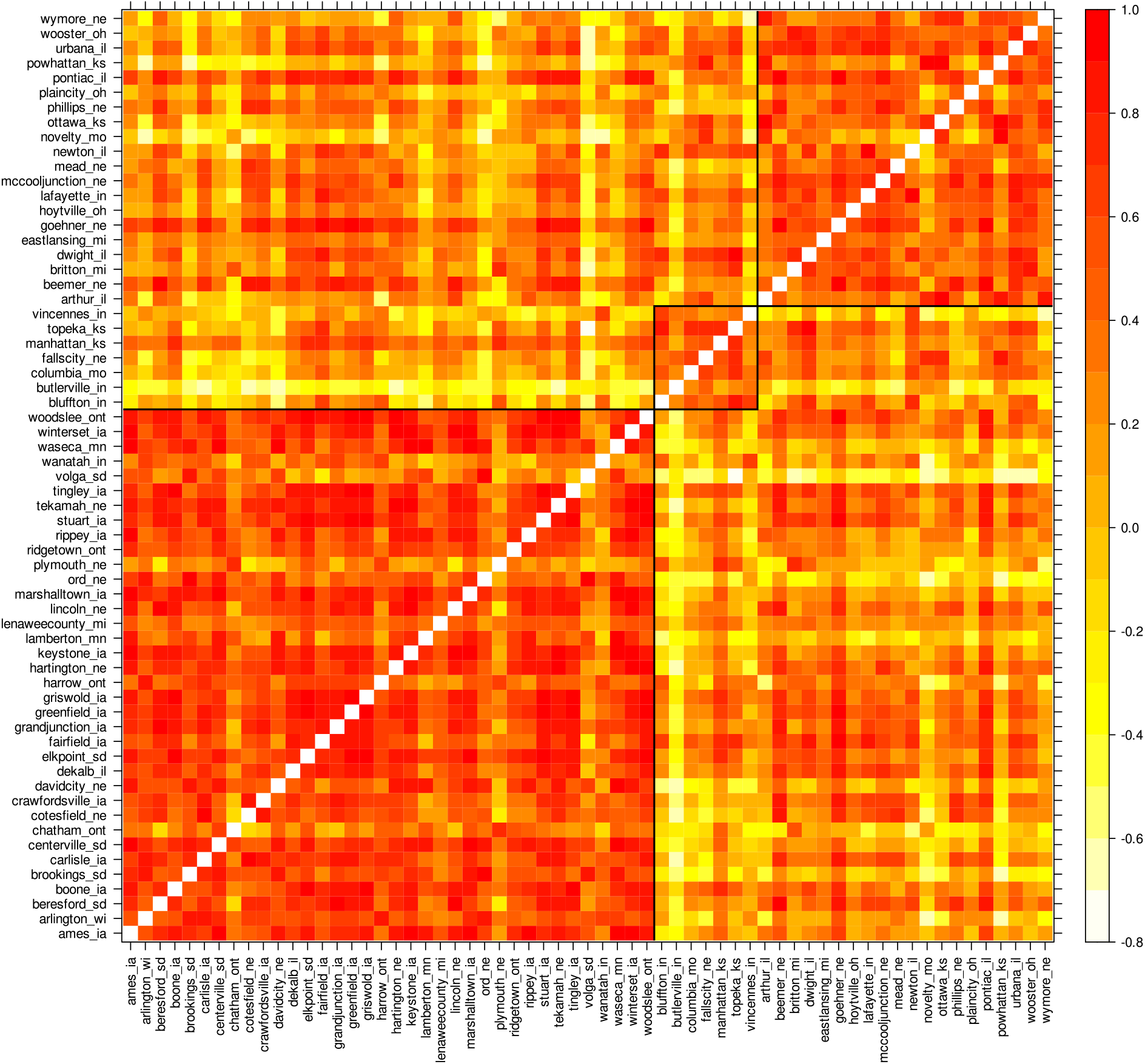
Heatmap of genetic correlations between the 63 observed locations from 1989 to 2019, estimated from the factor analytic (FA) model M2-18. Fitted K-means clusters are ordered from 1 to 3.

**Figure C6:**
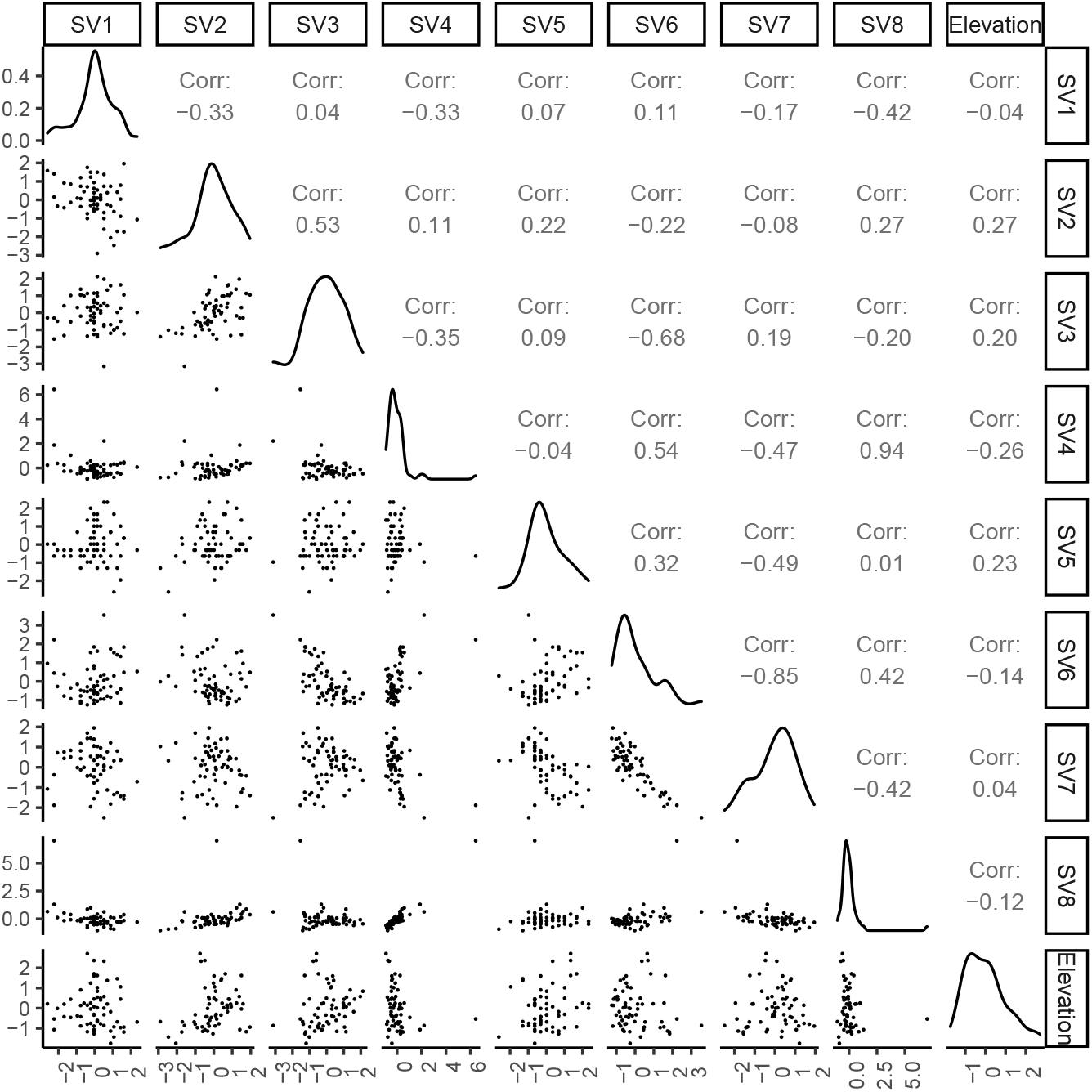
Scatterplots of the scaled and centered soil variables and elevation. Pearson correlation is displayed on the right. Variable distribution is available on the diagonal. The labels SV1, SV2, to SV8 are the soil variables described in section 5.3.5.

**Figure C7:**
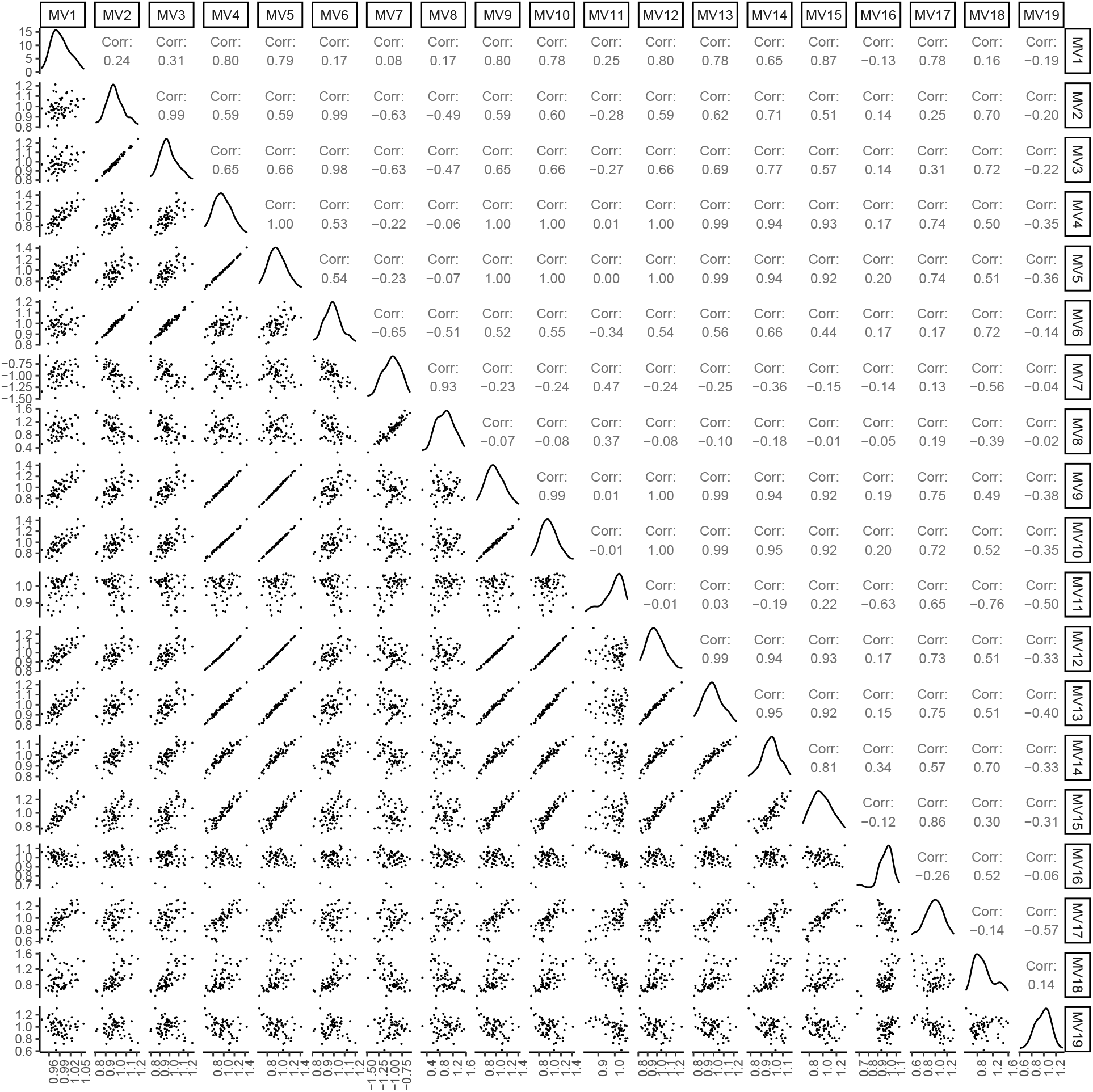
Scatterplots of the scaled and centered weather variables computed across years. Pearson correlation is displayed on the right. Variable distribution is available on the diagonal.The labels MV1, MV2, …, to MV19, are the weather variables described in Table C1.

**Figure C8:**
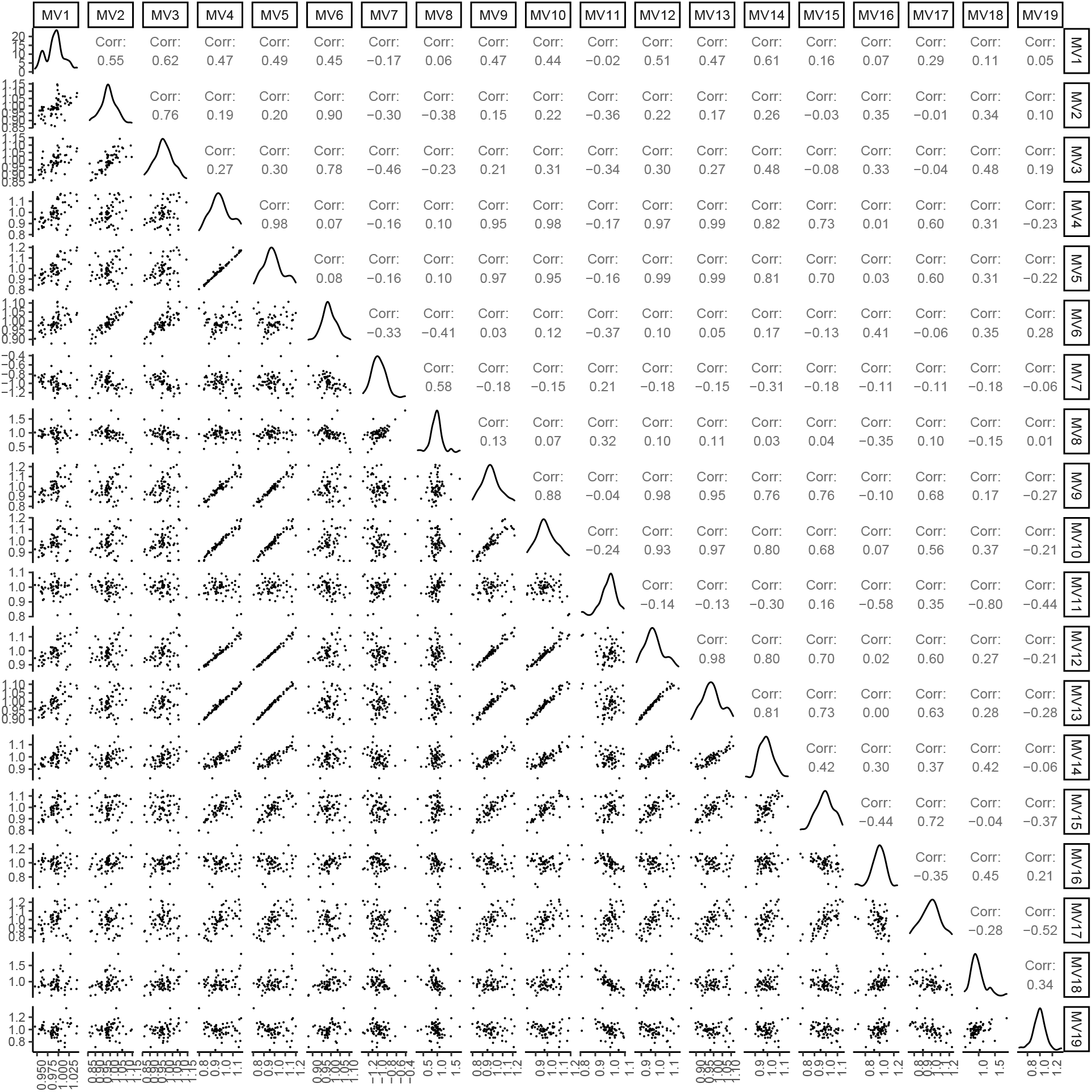
Scatterplots of the scaled and centered weather variables computed within years. Pearson correlation is displayed on the right. Variable distribution is available on the diagonal. The labels MV1, MV2, …, to MV19, are the weather variables described in Table C2.

**Figure C9:**
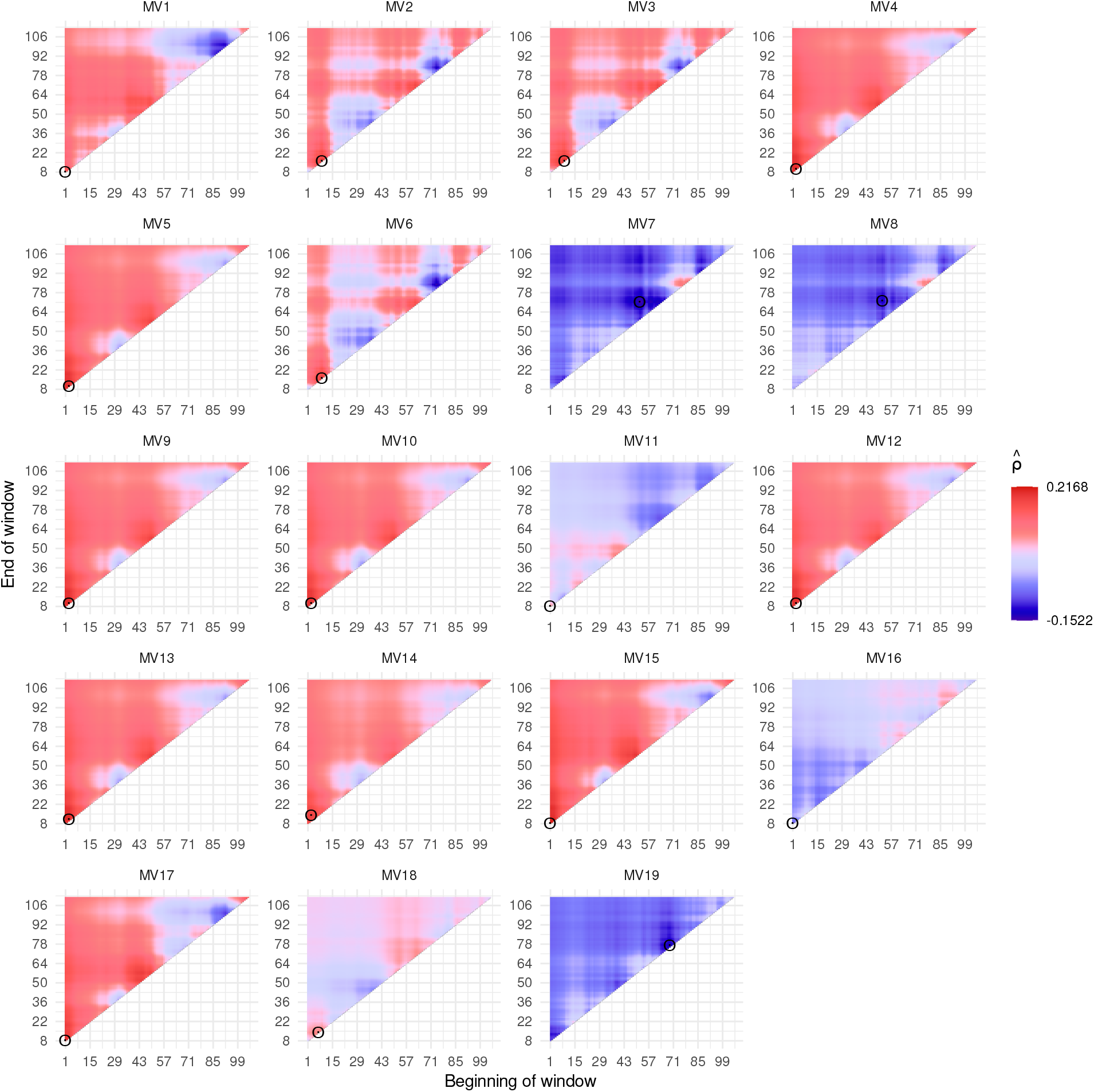
Exhaustive search from the critical environmental window computed from the Pearson correlation between the weather variables across years and the genotype by location deviations of each environment (location-year combination). The dots depict the highest window correlation. The labels MV1, MV2, to MV19 are the weather variables described in Table C1.

**Table C5:**
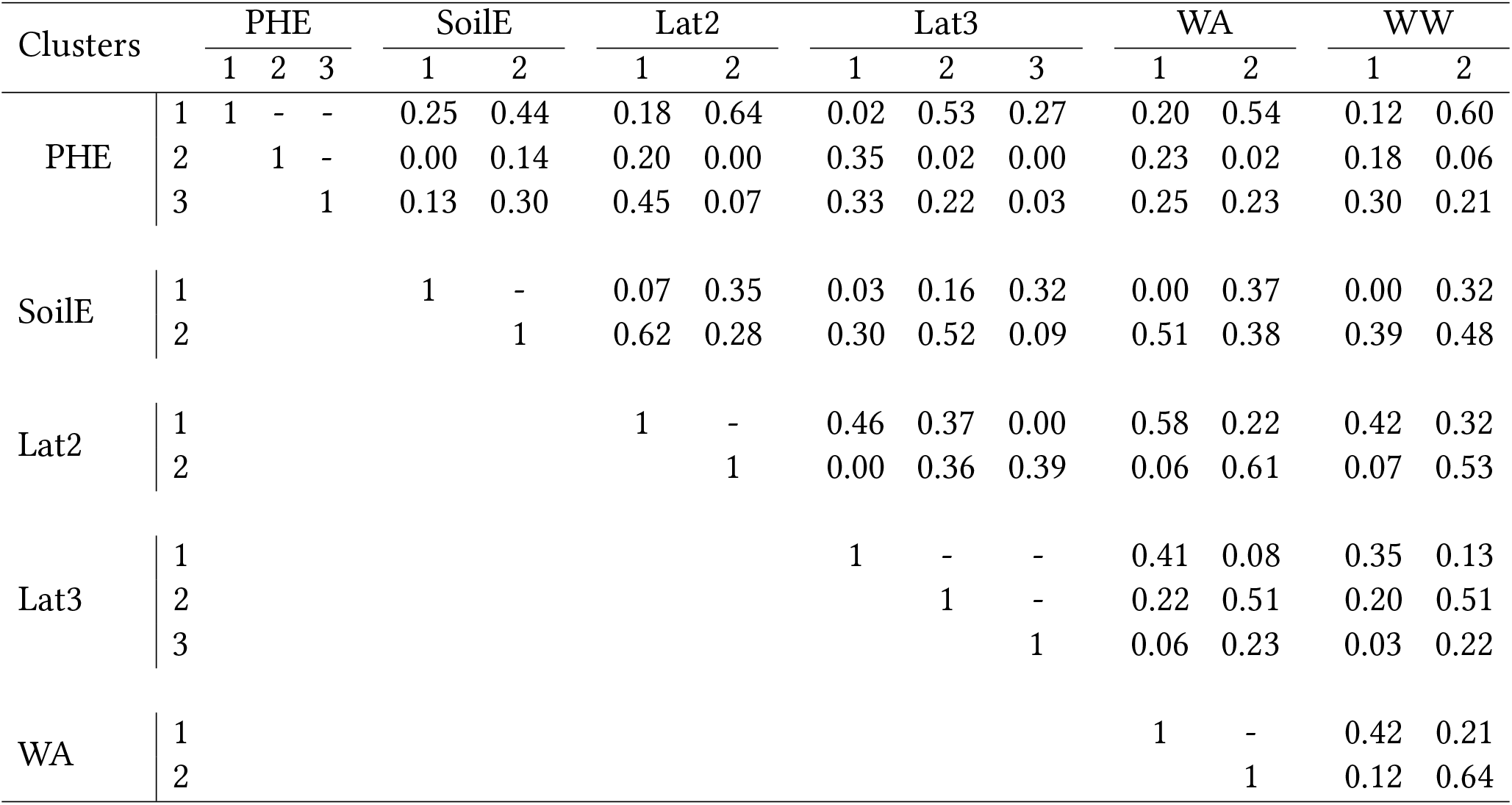
Jaccard similarity matrix between clustering-types (phenotypic - PHE, soil and elevation - SoilE, latitude split into 2 clusters - Lat2, latitude split into 3 clusters - Lat3, weather across years - WA, and weather within years - WW).

**Figure C10:**
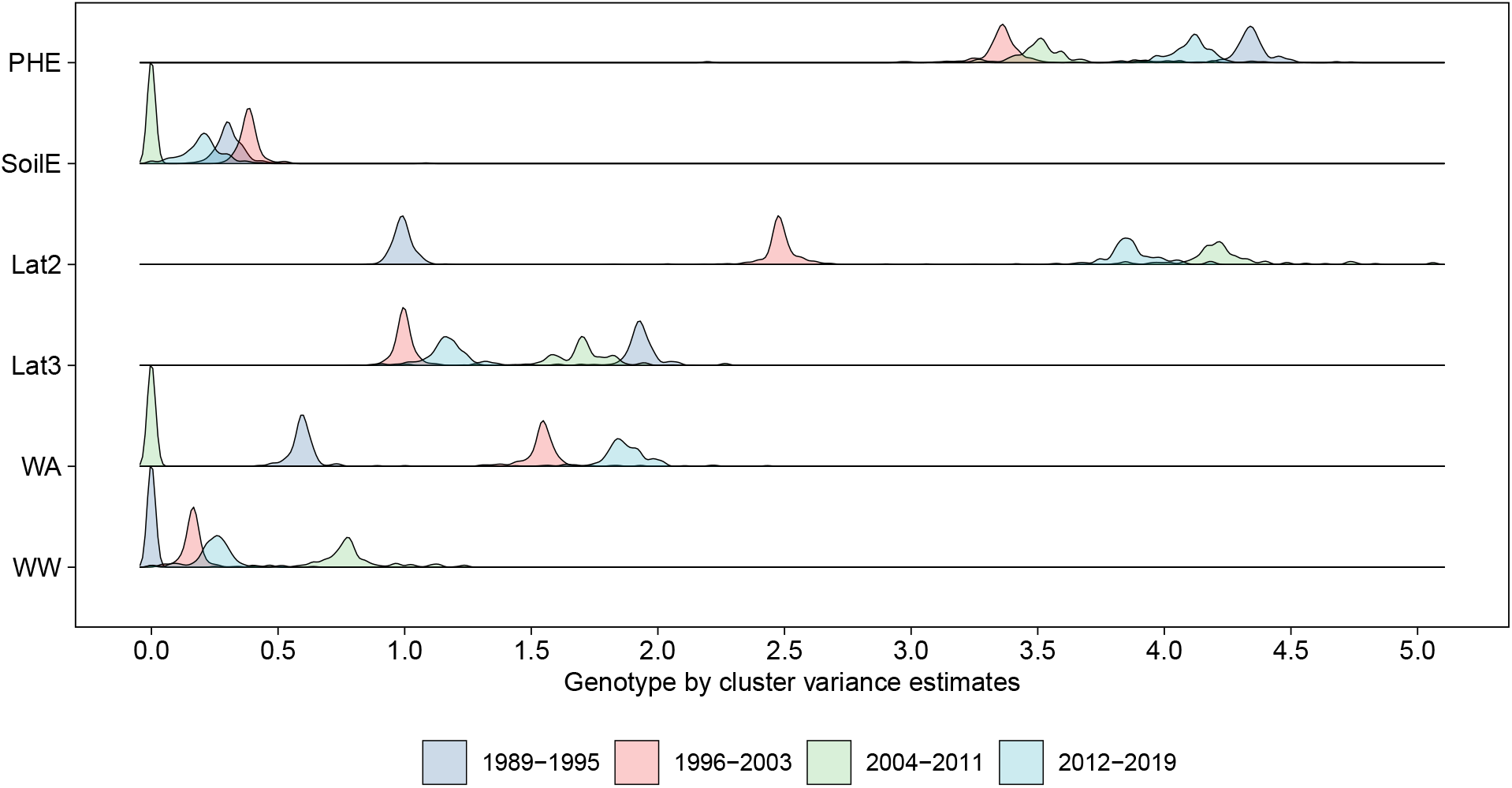
Jackknife estimates of genotype by cluster variances for the groups of years 1989- 1995, 1996-2003, 2004-2011, and 2012-2019, for phenotypic (PHE), soil + elevation (SoilE), latitude split into two groups (Lat2), latitude split into three groups (Lat3), weather across years (WA), and weather within years (WW) clustering types.

## 17 Appendix D - A variation of Model 3

Here we present a variation of Model 3 in which the genotype by environment interaction is included. This model was designed to be used with CERIS, where locations within years (i.e. environments) were represented as the average of the squared residuals (i.e., environmental means). The model is the following:

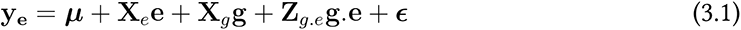

where **y_e_** is a vector of eBLUE values in a given year (i.e. **y_2_** in a given year), ***µ*** is the intercept, ***e*** is a vector of fixed effects of environments, ***g*** is a vector of fixed effects of genotypes, ***g.e*** is a vector of random effects of genotype by environment interaction with 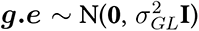, and ***E*** is a vector of residuals with ***E*** *∼* N(**0**, **Σ_e_**). **X***_e_*, **X***_g_*, and **Z***_g.e_* are incidence matrices for their respective effects, and **Σ_e_** was previously defined. With Model 3.1, locations within years were represented as the average of the observed predicted genotype by location values (i.e., eBLUP values). Then, the same procedure described in Sections 5.3.5 and 5.3.5.3 was applied.

The results are presented below, and in summary, are similar to the results from Table 4, Figures 5, and C4. The Jaccard similarity (data not shown) was also similar to Table C5. The 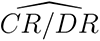 for WA using Model 3 was 0.92 (Table 4), which is slightly superior to the value obtained using Model 3.1 (Table D1).

**Table D1:** The ratio of correlated responses from selection across all environments relative to direct responses to selection within mega-environments (CR/DR) for weather variables using Model 3.1. *ρ_g_* is the correlation between estimated genotypic effects in the non-clustered and clustered sets of environments, 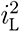 and 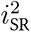 are the reliabilities of genotype means in the non-clustered and clustered sets of environments, respectively. Weather from means across years (WA), and weather from means within years (WW).

| Clustering type<sup>a</sup> | Number of |  | Estimates of |  |  |  |
| --- | --- | --- | --- | --- | --- | --- |
| | Clusters | Locations | $\hat{\rho}_g$ | $\hat{i}_L^2$ | $\hat{i}_{SR}^2$ | $\widehat{CR/DR}$ |
| WA | 2 | 23 / 40 | 0.94 | 0.48 | 0.45 | 0.98 |
| WW | 2 | 30 / 33 | 0.97 | 0.49 | 0.46 | 1.00 |

**Figure D1:**
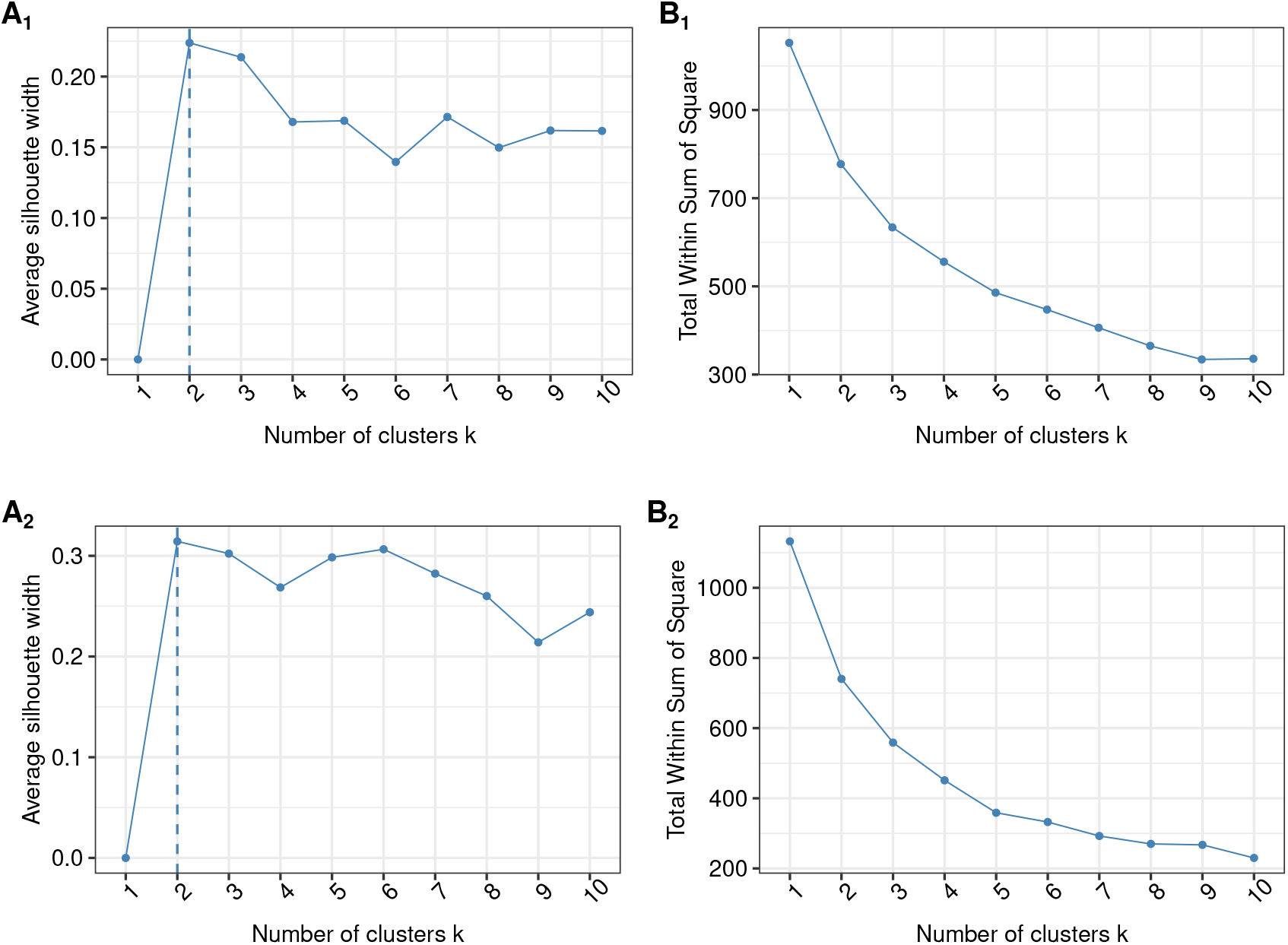
Graphical display of the optimal number of clusters based on the Silhouette (A) and Elbow (B) methods for weather within year variables (WW, A_1_ and B_1_) and weather across year variables (WA, A_2_ and B_2_). CERIS was implemented with Model 3.1.

**Figure D2:**
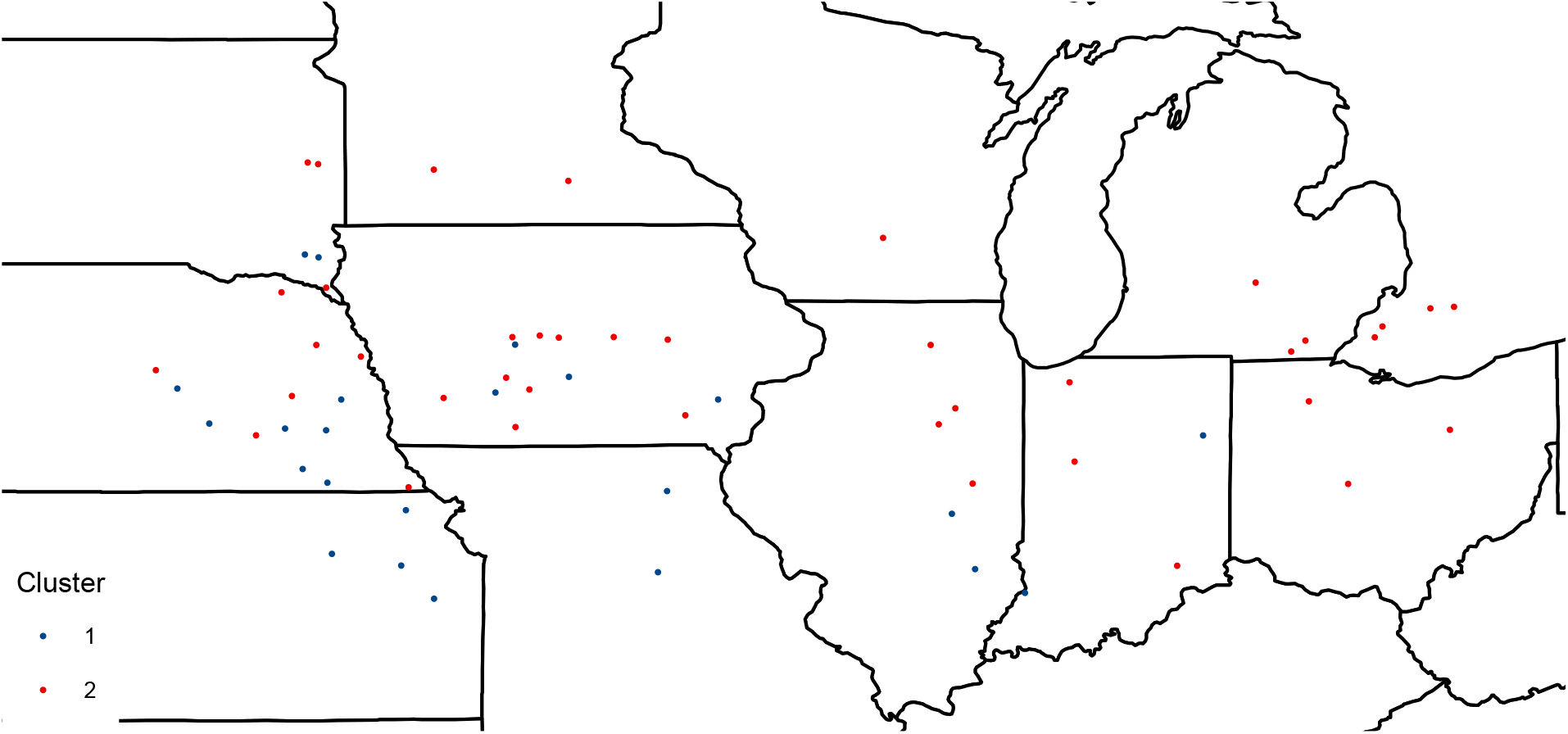
Geographic visualization of the target population of environments divided according to weather across years (E), computed with Model 3.1.

